# Mutation in *Ralstonia pseudosolanacearum* F1C1 cytochrome *p450* homologue impairs virulence through root but not through shoot of tomato plants

**DOI:** 10.64898/2026.10.09.757812

**Authors:** Lukapriya Dutta, Kristi Kabyashree, Shubhra Jyoti Giri, Kanika Bansal, Disha Chakraborty, Shuhada Begum, Sumi Kalita, Monika Jain, Shuvam Bhuyan, Sunita Kushwah, Aditya Kumar, Gopaljee Jha, Prabhu B. Patil, Suvendra Kumar Ray

## Abstract

*Ralstonia pseudosolanacearum* causes a lethal bacterial wilt disease in many plants. The pathogen enters through roots and systemically colonizes the xylem tissue, resulting in wilting and eventual death of the host. Using transposon mutagenesis followed by screening in tomato seedlings, we identified a mutant exhibiting virulence deficiency by root inoculation, but not by leaf inoculation. The mutation was recreated in the wild-type genome via natural transformation to prove the linkage of virulence deficiency with the transposon insertion. Whole genome sequencing revealed that the transposon insertion was in a *cyp450* homologue, encoding cytochrome P450 monooxygenase. Two independent targeted mutations were generated in the *cyp450* gene, and both the mutants recapitulated the root-specific virulence deficiency. The mutants rarely cause disease in mature plants following soil drenching but remained fully virulent upon petiole pricking inoculation. Complementation with the full length *cyp450* gene amplified from the wild-type strain F1C1 restored virulence in the mutant. Compared to the wild-type F1C1, the *cyp450* mutants exhibited greater sensitivity to tomato root extracts. HR-LCMS of tomato root extracts revealed the occurrence of several antimicrobial compounds. Cytochrome P450 enzymes are known to mediate the degradation or biosynthesis of diverse metabolites in bacteria. Considering the virulence difference due to the mode of inoculation, our findings indicate that *cyp450* contributes to tissue-specific adaptation of the pathogen during infection has opened an avenue for integrating tissue metabolomics with bacterial wilt susceptibility research.

## 1 Introduction

Bacterial wilt, caused by the strains of *Ralstonia solanacearum* species complex (RSSC), is one of the most devastating plant diseases that poses an enormous risk to agricultural productivity worldwide (Saini et al. 2025a). The pathogen has a wide host range, infecting more than 500 plant species across 86 botanical families (Lowe-Power et al. 2020). Among RSSC members, *R. pseudosolanacearum* comprises phylotype I and phylotype III, significantly affects solanaceous crop plants like tomato, potato, eggplant, chilli, causing severe yield and quality losses in tropical and subtropical regions of Asia and Africa, respectively (Saini et al. 2025b; Vailleau and Genin 2023). Major virulence determinants of this bacterium include effectors secreted by type III secretion systems, cell wall-degrading enzymes secreted by type II secretion systems, and bacterial motility associated traits (Buttner et al. 2009; Kang et al. 1994; Tans-Kersten et al. 2001; Vailleau and Genin 2023).

The pathogen survives for extended periods in soil and water until it senses the roots of the host plant and penetrates the host at the sites of root emergence or wounds (de Pedro-Jové et al. 2021; Yao et al. 2006). Root plays a crucial role during plant colonization as the pathogen initially attaches on roots following entry through sites of lateral root emergence (Caldwell et al. 2017; Vailleau and Genin 2023). Subsequent systemic colonization in the host, together with obstruction of the vascular bundles by excessive secretion of exopolysaccharides lead to wilting (Genin 2010; Planas-Marquès et al. 2020). Although known as a xylem pathogen, it has to adapt distinct tissue physiologies during different stages of infection. Therefore, identifying tissue- or host-specific virulence factors may provide novel insight into pathogen adaptation inside the host. Despite extensive studies on virulence factors, reports concerning tissue-specific virulence factors of this systemic phytopathogen remain limited. Previous reports have revealed that HrpB, and HrpG deficient strains differentially affect root growth in *Arabidopsis thaliana* during pathogenesis (Lu et al. 2018; Zhang et al. 2024). Research on pathogen colonization patterns in root and shoot tissues of resistant and susceptible tomato varieties demonstrated an impaired systemic colonization of the pathogen in resistant varieties (Planas-Marquès et al. 2020). *R. pseudosolanacearum* F1C1 (Kumar et al. 2013) has been demonstrated to cause wilting in tomato seedlings through inoculation either in root or in cotyledon leaves, with a higher magnitude of virulence in the latter (Kabyashree et al. 2020; Kumar et al. 2017; Sing et al. 2018a). Brinjal seedlings exhibit a greater susceptibility than tomato seedlings upon leaf inoculation compared to tomato, whereas the reverse trend is observed following root inoculation (Bhuyan et al. 2026, 2025; Phukan 2018). Collectively, these observations indicate that certain virulence factors might function in a tissue-specific manner during colonization and infection (Bhuyan et al. 2026, 2025; Kabyashree 2021). Therefore, exploring tissue-specific virulence determinants of the pathogen is both timely and necessary.

Although canonical virulence determinants in RSSC are well characterized, the genetic basis of tissue-specific adaptation remains largely unexplored. In the present study, we are using random insertion mutagenesis coupled with genomics to explore role of metabolic adaptation in shaping pathogen fitness across distinct host tissues. Here we have employed a forward genetics approach to generate transposon-induced mutants of *R. pseudosolanacearum*. Screening for the virulence phenotype in tomato seedlings led to the identification of a mutant that exhibited virulence deficiency upon root inoculation but not upon leaf inoculation. Whole genome sequencing along-with molecular genetics approach confirmed the role of *cyp450* gene in mediating differential virulence. To our knowledge, this study provides the first evidence implicating *cyp450* gene in virulence of any phytopathogenic bacterium. In phytopathogenic bacteria, the gene may be required for the adaptation to the plant environment.

## 2 Results

### 2.1 Transposon-induced mutants exhibit virulence deficiency by root inoculation but not by leaf inoculation in tomato seedlings

*R. pseudosolanacearum* F1C1 is virulence proficient in tomato seedlings upon either leaf or root inoculations. The transposon mTn5SSgusA11-induced F1C1 mutants were generated by executing biparental conjugation between TRS1001 (Rif^r^ F1C1) as the recipient and *Escherichia coli* TP003 as the donor carrying the transposon on a suicide plasmid. The mTn5 induced mutants were screened for their virulence in tomato seedlings by both root and leaf inoculations. Out of 140 mutants screened, TRS2005, TRS2006, and TRS2007 displayed virulence deficiency upon root inoculation but retained virulence upon leaf inoculation (Kabyashree 2021). The TRS2005 mutant was selected for further characterization. To prove that the virulence phenotype of TRS2005 was linked to the mTn5 insertion, the mutation was recreated in the wild-type F1C1 background via natural transformation using the genomic DNA from TRS2005. Two independent transformants TRS2005.1 and TRS2005.2 were obtained and evaluated for pathogenicity in tomato seedlings following both root and leaf inoculations. Both exhibited virulence phenotype similar to that of the parent mutant, confirming the linkage of the mTn5 insertion and altered virulence phenotype (Figure S1).

Whole genome sequencing of the TRS2005 mutant was carried out to identify the insertion site of mTn5. Illumina paired-end sequencing libraries (2 x 250 read length) from genomic DNA, and *de novo* assembly identified single insertion in the genome at cytochrome p450 monooxygenase reductase (*cyp450*). The gene is present in a monocistronic operon in the F1C1 genome (http://14.139.219.242:8009/), and its ORF size is 1197 bp that encodes a protein of 398 amino acids (Kabyashree 2021).

### 2.2 *cyp450* mutant creation and complementation with the full length *cyp450* gene

To study the role of the *cyp450* gene in the virulence of F1C1, two targeted mutations were created in the *cyp450* gene (described in Figure S2a and S2b). Creation of two independent *cyp450* mutants *cyp450*::*lacZ* (TRS5001), *cyp450*::ΩSp (TRS5002) and complementation with full length *cyp450* gene, *cyp450*::ΩSp::*cyp450*-COMP (TRS5003), has been described in details in the Materials and Methods. The insertion mutation was confirmed by PCR amplification in the *cyp450*::*lacZ* (TRS5001) genome. No amplification was detected in the wild type F1C1 genome (Figure 1a). The mutant *cyp450*::*lacZ* (TRS5001) is positive for *ß*-galactosidase activity. The expression of the *cyp450* gene was accessed by studying *β*-galactosidase activity which shows similar expression of *cyp450* gene in BG as well as in minimal medium (Figure 1d). The insertion-deletion mutant *cyp450*::ΩSp (TRS5002) was confirmed by a PCR amplification (Figure 1b). The restoration of full length *cyp450* gene in *cyp450*::ΩSp::*cyp450*-COMP (TRS5003) genome was confirmed by PCR amplification (Figure 1c) The complemented strain is positive for *ß*-Galactosidase activity in BG as well as in minimal medium (Figure 1d).

**FIGURE 1.**
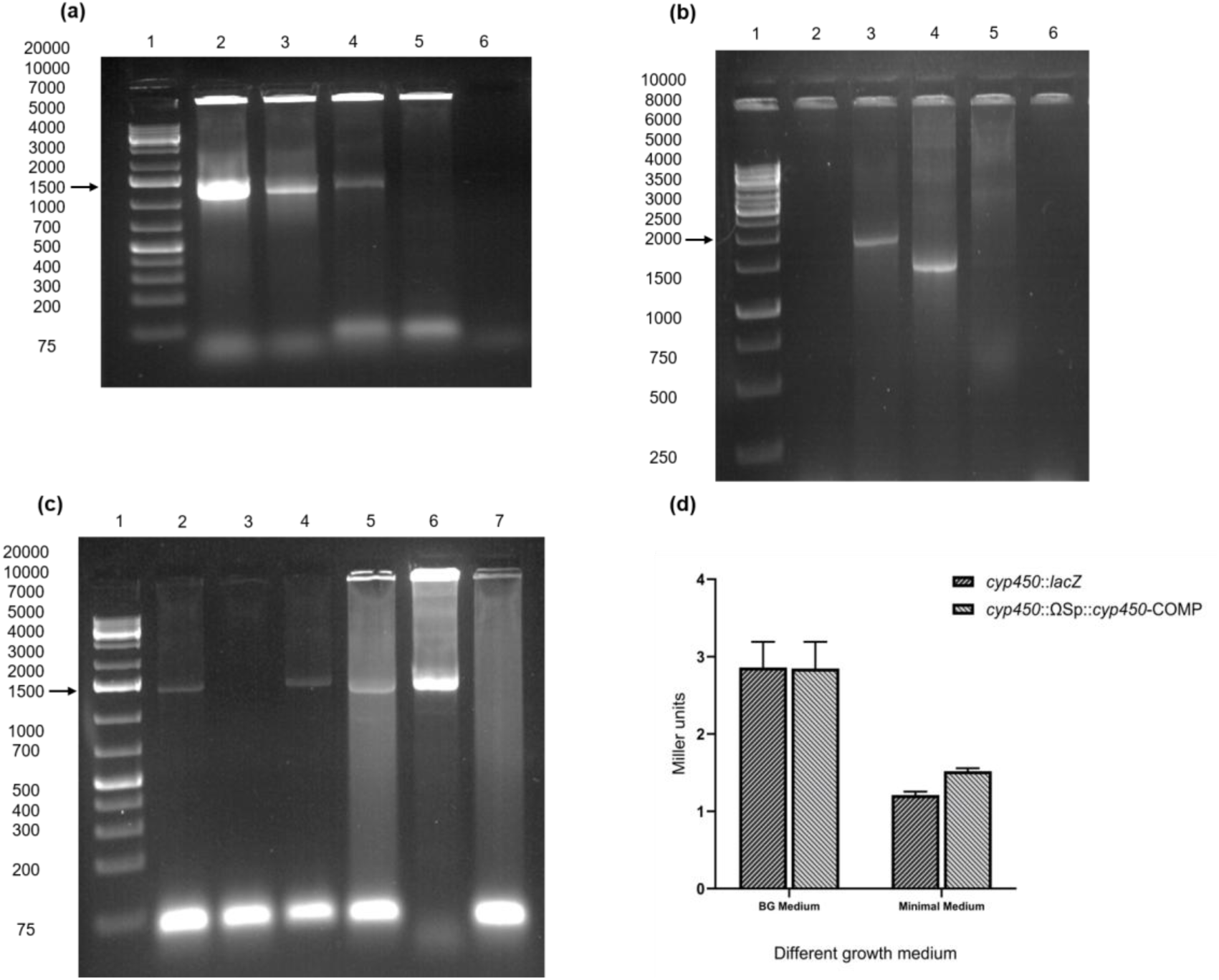
Creation and confirmation of *cyp450* insertion mutant, insertion-deletion mutant and complemented strain. (a) In this figure, Lane 1 has 1 kb plus DNA ladder. Lane 2, 3 and 4 shows a desired amplification of ∼1.3 kb (1287 bp) in *cyp450*::*lacZ* (TRS5001) using oligo against the lacZ and oligo against the upstream region of the amplicon. It proved the insertion in the *cyp450* gene. No such amplification was observed in lane 5, wild type F1C1 genome as there was no *lacZ* gene. Lane 6 is the control having no amplification as no template added. (b) In this figure, Lane 1 has 1 kb DNA ladder. Lane 2 having no amplification in mutant *cyp450*::ΩSp (TRS5002). Lane 3 showing desired amplification of size ∼ 2 kb (1904 bp) in wild type F1C1. Lane 4 showing desired amplification of size ∼ 1.4 kb (1397 bp) in plasmid pLD005 carrying the 648 bp fragment I and 749 bp fragment II (deletion of 507 bp). Lane 5 has no amplification in the plasmid pLD006 carrying ΩSp insertion between fragment I and fragment II. Lane 6 is the control having no template. (c) In this figure, Lane 1 has 1 kb plus DNA ladder. Lane 2 showing desired amplification of ∼ 1.4 kb (1346 bp) in the wild type F1C1. Lane 3 showing no amplification in mutant strain *cyp450*::ΩSp (TRS5002). Lane 4 and 5 showing desired amplification of ∼ 1.4 kb (1346 bp) in complemented strain *cyp450*::ΩSp::*cyp450*-COMP (TRS5003). Lane 6 showing desired amplification of ∼ 1.4 kb (1346 bp) in plasmid pLD007 carrying the complementation construct. Lane 7 is the control having no template control. (d) *β*-Galactosidase activity by a reporter gene fusion was performed in BG and minimal growth medium for the insertion mutant *cyp450*::*lacZ* (TRS5001) and complemented strain *cyp450*::ΩSp::*cyp450*-COMP (TRS5003). Similar gene expression was observed for both the medium. Error bar represents SD (n=3).

### 2.3 *cyp450* mutants are virulence deficient through root inoculation but proficient through shoot inoculation in tomato plants

The virulence phenotype of the mutants *cyp450*::*lacZ* (TRS5001), *cyp450*::ΩSp (TRS5002) was studied in tomato seedlings by inoculation in both root as well as leaves. Both the mutants exhibited virulence deficiency through root but not by leaf clip inoculation. After creating the complemented strain, virulence in tomato seedlings was studied with wild type F1C1, the two *cyp450* mutants *cyp450*::*lacZ* (TRS5001), *cyp450*::ΩSp (TRS5002), and the complemented strain *cyp450*::ΩSp::*cyp450*-COMP (TRS5003). In a single inoculation study, two sets of independent experiments were carried out where 40 seedlings per strain was inoculated per set. Total three inoculation studies were carried out. Wilting of the seedlings were scored from one dpi onwards till 10 dpi. In case of root inoculation, the wilting percentage of seedlings inoculated with wild type F1C1 was 48.75%, for *cyp450*::*lacZ* (TRS5001) the wilting percentage was 22.15%, for *cyp450*::ΩSp (TRS5002) the wilting percentage was 28.75%. The wilting percentage of the seedlings inoculated with complemented strain *cyp450*::ΩSp::*cyp450*-COMP (TRS5003) was 47.5%. The survival probability of the seedlings inoculated with wild type F1C1 was 45.95%, for the *cyp450*::*lacZ* (TRS5001) and *cyp450*::ΩSp (TRS5002) mutants the survival probability was 75.67% and 67.56%, respectively. For the seedlings inoculated with complemented strain *cyp450*::ΩSp::*cyp450*-COMP (TRS5003), the survival probability was 47.64% (Figure S4a). There was a distinct virulence deficiency in the two mutants in comparison to the wild type upon root inoculation. The complemented strain virulence was significantly higher than both the mutants. In case of leaf inoculation in the seedlings, the virulence of the mutants *cyp450*::*lacZ* (TRS5001), and *cyp450*::ΩSp (TRS5002) were equally virulent like the wild type F1C1 and complemented strain *cyp450*::ΩSp::*cyp450*-COMP (TRS5003). In case of the wild type F1C1, the wilting percentage was 80%, for the mutants *cyp450*::*lacZ* (TRS5001) and *cyp450*::ΩSp (TRS5002) the wilting percentage was 76.25% and 81.25%, respectively. For the complemented strain *cyp450*::ΩSp::*cyp450*-COMP (TRS5003), the wilting percentage was 100%. The survival probability was also calculated. The survival probability of the seedling inoculated with wild type F1C1 was 11.26%, for the mutants *cyp450*::*lacZ* (TRS5001) and *cyp450*::ΩSp (TRS5002) the survival probability was 18.53% and 10.13%, respectively. For the complemented strain *cyp450*::ΩSp::*cyp450*-COMP (TRS5003), the survival probability was 4.76% (Figure S4b). The wilting score was higher in the case of leaf inoculation than that in the case of root inoculation.

The virulence phenotype of the *cyp450*mutants was studied in the grown-up plants by inoculation in the root following soil drenching method. Two independent pathogenicity experiments were conducted. In the first experiment of soil drenching, wilting symptoms were first observed at 4 dpi in two plants inoculated with the complemented strain *cyp450*::ΩSp::*cyp450*-COMP (TRS5003). By the end of the experiment 11 plants inoculated with F1C1 and 13 plants inoculated with *cyp450*::ΩSp::*cyp450*-COMP (TRS5003) wilted, compared with 3 plants inoculated with *cyp450*::*lacZ* (TRS5001) (p<0.05) and 1 plant inoculated with *cyp450*::ΩSp (TRS5002) (p<0.05) wilted, whereas no mortality was observed in the plants inoculated with sterile distilled water (Figure 2a). The random occurrence of disease was represented as color filled circles (Figure 2b). Disease scores recorded at each dpi were used to generate a disease progression curve, illustrating wilting symptoms developed over time (Figure 2c). Similar observations were obtained in the second independent experiment of soil drenching and the data are presented in Figure S5.

**FIGURE 2.**
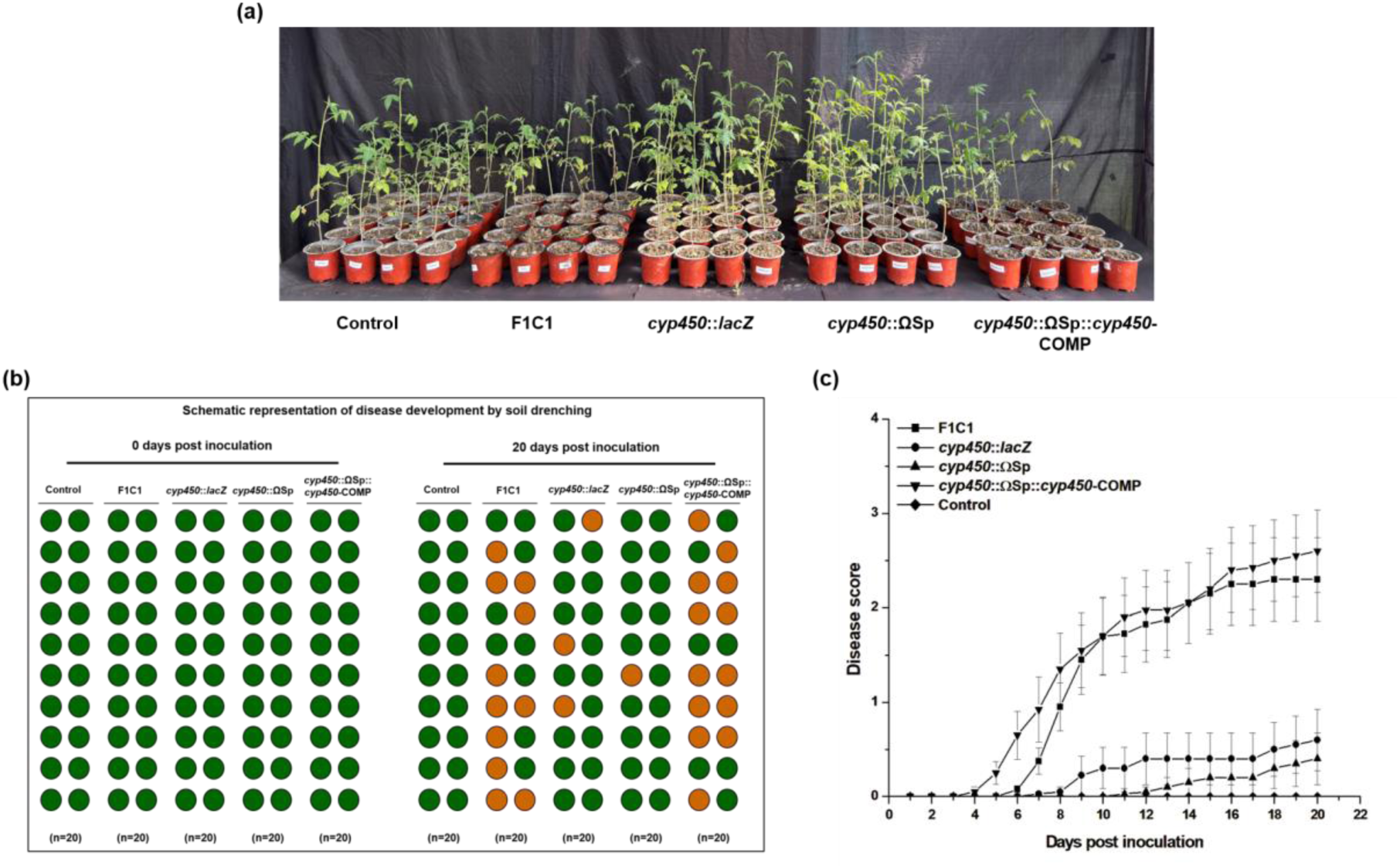
Virulence phenotype of *cyp450* mutants in grown-up tomato plants by soil drenching. In Fig. (a) 20 plants per strain were inoculated by soil drenching method. 50 ml of the respective bacterial inoculum (OD_600_=0.6) per plant was poured into the soil in each pot. The photograph was taken at 20 dpi. 11 plants wilted in case of wild type F1C1, 13 plants wilted in case of complemented strain *cyp450*::ΩSp::*cyp450*-COMP (TRS5003). 3 plants wilted in case of *cyp450*::*lacZ* (TRS5001) and 1 plant wilted in case of *cyp450*::ΩSp (TRS5002) representing significant reduction in virulence compared to wild type and complemented strain. The all 20 plants in the control were healthy. For easy visualization of the disease, the pots were arranged properly in case (a), The random occurrence of the disease as well as the escapees among the inoculated plants are schematically presented in figure (b) as shown by filled circle, healthy plants in green and wilted plants in brown. The disease progression is represented in Figure (c). Plants were monitored regularly and wilting symptoms were scored till 20 dpi at a disease index scale from 0-4. The graph represents one independent experiment consisting 20 plants per strain.

However, this experiment was performed earlier with a sample size (n=5) for each strain, with three independent experiments. So, in total, five times this experiment was performed by soil drenching, and every time virulence deficiency of the two *cyp450* mutants was observed in comparison to the wild-type F1C1.

To study the disease phenotype through shoot of grown-up plants, bacteria was directly inoculated in the petiole of the plant. Two independent petiole pricking experiments were conducted and disease scores were recorded up to 15 dpi. In the first experiment, wilting symptoms of the compound leaves were first observed at 2 dpi, and disease progression was monitored and scored until 15 dpi. By the end of the experiment, all 12 plants inoculated independently with F1C1, *cyp450*::*lacZ* (TRS5001), *cyp450*::ΩSp (TRS5002), and *cyp450*::ΩSp::*cyp450*-COMP (TRS5003) were wilted. No mortality was observed in the water control group Figure 3a. The disease development is schematically presented by the color filled circles Figure 3b. Disease scores recorded throughout the 15 days were used to construct a disease progression curve Figure 3c. Similar observations were obtained in the second independent experiment of petiole pricking and the data are presented in supplementary Figure S6.

**FIGURE 3.**
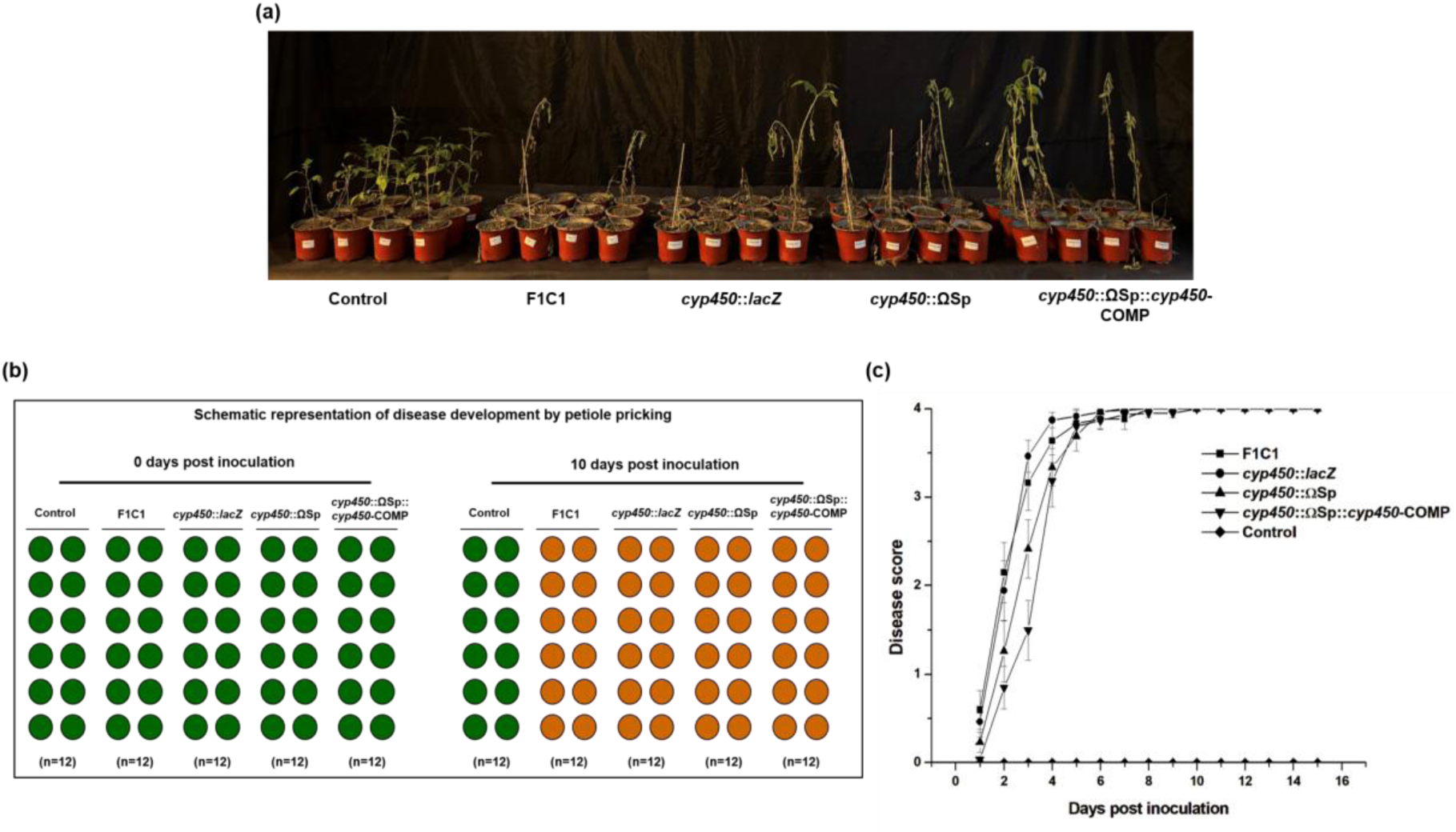
Virulence phenotype of *cyp450* mutants in grown-up tomato plants by petiole pricking. (a) 12 plants per strain such as wild-type F1C1, *cyp450*::*lacZ* (TRS5001), *cyp450*::ΩSp (TRS5002), *cyp450*::ΩSp::*cyp450*-COMP (TRS5003) (∼10^9^ CFU/ml) and water control were inoculated by petiole pricking method. The photograph was taken at 15 dpi. All the 12 plants inoculated with the bacterial strains developed wilting. Plants inoculated with sterile distilled water remained healthy throughout the experiment. The disease development is schematically presented by the color filled circles as greens represents healthy plants and brown represents wilted plants in Figure (b). The disease progression is represented disease in Figure (c).

However, this experiment was performed earlier with a sample size (n=3) for each strain, with three independent experiments. So, in total five times this experiment was performed in grown up plants. Every time similar virulence proficiency of the two *cyp450* mutants was observed like wild type F1C1 in petiole pricking.

### 2.4 Root extract of tomato seedlings inhibits the growth of *cyp450* mutants and it contains several antibacterial compounds

The impact of the root as well as the leaf extracts of tomato seedlings on the growth of *cyp450* mutants, complemented strain as well as the wild type F1C1 was studied qualitatively on BG agar medium. Leaf extract had no visible inhibitory impact on the growth of any of the strains such as *cyp450* mutants, complemented strain and the wild type F1C1 (Figure 4d, e and f). However, the root extract had a differential growth inhibition effect on the *cyp450* mutants in comparison to the wild type F1C1 and complemented strain and wild type F1C1. In the plate assay a distinct inhibition of the growth of the mutants such as *cyp450*::*lacZ* (TRS5001) and *cyp450*::ΩSp (TRS5002), was observed near to the wells containing tomato root extract in comparison to the growth of the wild type F1C1 and the complemented strain (Figure 4a, b and c).

**FIGURE 4.**
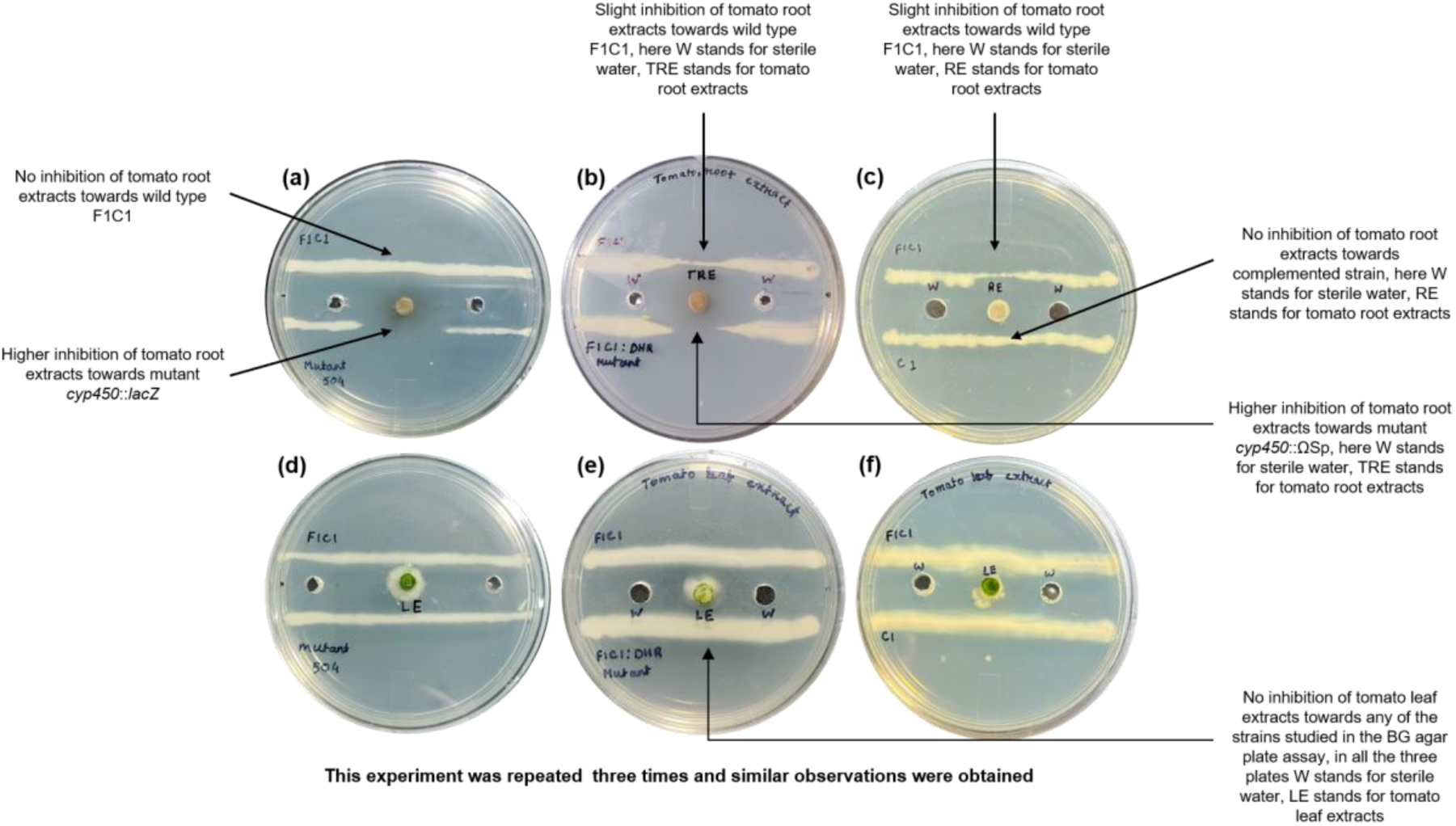
Tomato seedlings root and leaf extracts sensitivity assay. Study of bacterial growth regarding inhibition of tomato root extracts and tomato leaf extracts. In BG agar medium three wells of 4.0 mm diameter were made. One well positioned at the center of the plate and the other two wells located 2 cm away from the central well. Wild type F1C1 was streaked close to the well at an equal distance of 1 cm above. The mutants and complemented strain were streaked close to the well at an equal distance of 1 cm below. 50 µL of the tomato root and leaf extract was added to the middle well, 50 µl of the sterile distilled water was added which was taken from the distilled water washing step of surface sterilization. (a) Higher growth inhibition of tomato root extracts was observed towards insertion mutant *cyp450*::*lacZ* (TRS5001) but not towards wild type F1C1. (b) Higher growth inhibition was observed towards insertion-deletion mutant *cyp450*::ΩSp (TRS5002) and a slight growth inhibition was observed towards wild type F1C1 (c) No growth inhibition was observed towards complemented strain *cyp450*::ΩSp::cyp450-COMP (TRS5003). (d) (E) (F) No inhibitory effect of tomato leaf extracts was observed towards any of the strains such as, wild type F1C1, *cyp450* mutants *cyp450*::*lacZ* (TRS5001), *cyp450*::ΩSp (TRS5002) and complemented strain *cyp450*::ΩSp::*cyp450*-COMP (TRS5003).

The experiment was repeated for three times and the inhibition of *cyp450* mutants was consistently observed in all the three independent experiments. Additionally, an agar well diffusion assay was performed to evaluate the sensitivity of the strains towards tomato root extract by measuring the zone of inhibition. Observation was recorded up to 48 h. The mutants *cyp450*::*lacZ* (TRS5001) and *cyp450*::ΩSp (TRS5002) exhibited a higher zone of inhibition in comparison to the wild type F1C1 and the complemented strain *cyp450*::ΩSp::*cyp450*-COMP (TRS5003), as represented in (Figure S8). The higher sensitiveness of the mutants than the wild type F1C1 and the complemented strain towards the root extract was in concordance with the observation of virulence deficiency through root inoculation.

The inhibitory effect of the root extracts against the growth of *cyp450* mutants indicated the presence of antibacterial compounds. Accordingly, the HR-LCMS profiling identified several phytochemicals in methanolic extracts of tomato seedling roots. Numerous phytoconstituents were identified in the root extract, of which 20 were detected for exhibiting antibacterial properties (Table 4). The compounds identified in the extracts were mainly phenolic, alkaloid, and steroid/terpenoid groups. Sensitiveness of the *cyp450* mutants specific to any of these antibacterial compounds will be studied in future.

### 2.5 *cyp450* mutants presence in tomato seedlings inoculated through root

The occurrence of the mutants *cyp450*::*lacZ* (TRS5001), *cyp450*::ΩSp (TRS5002), F1C1, and complemented strain *cyp450*::ΩSp::*cyp450*-COMP(TRS5003) in tomato seedlings were studied after root inoculation. At 4 dpi, the occurrence of F1C1, *cyp450*::*lacZ* (TRS5001), *cyp450*::ΩSp (TRS5002) and *cyp450*::ΩSp::*cyp450*-COMP (TRS5003) was found out by crushing the seedlings and dilution plating the extract on BG-agar plates. The presence of the wild type and the mutants were observed in the inoculated seedlings (Table S2). To observe it more critically, a mixed-inoculation experiment between the wild type and the mutants was performed in seedlings. At 0, 2 and 4 dpi, both mutants and wild type strains were isolated from the inoculated seedlings in all the cases. The increase in the number of the wild type and the mutants was observed in the seedlings after inoculation (Table S1). The isolation of the mutants from the inoculated seedlings were obvious considering the occurrence of disease by the mutant though at a lesser magnitude. This study was based on bacterial population in the entire seedling. A tissue specific population level comparison between the wild type F1C1 and the mutant is required to understand the mutant population in the root and the leaves. Similarly, the occurrence of the mutants in inoculated seedlings were also observed by inoculating GFP tagged mutant (*cyp450*::*lacZ*/pDSK-GFPuv) (Figure S9).

### 2.6 *cyp450* mutants exhibit similar growth rate, extracellular enzyme activity, motility, hypersensitive response to the wild type F1C1

The *cyp450* insertion mutant *cyp450*::*lacZ* (TRS5001) and insertion-deletion mutant *cyp450*::ΩSp (TRS5002), along with the complemented strain *cyp450*::ΩSp::*cyp450*-COMP (TRS5003), were evaluated for generation time, production of extracellular enzymes such as cellulase and pectinase, type-IV pili mediated twitching motility, flagellar motility and hypersensitive response (Figure S10). Mutants were indifferent from the wild type F1C1 with regard to the above characteristics, which indicates that the virulence deficiency in the mutants was not due to deficient in any of these characteristics.

## 3 Discussion

In this study we have used molecular genetic approaches to characterize *R. pseudosolanacearum* virulence functions, which are required during infection through root but not through shoot. Transposon induced *R. pseudosolanacearum* mutants have been screened for virulence both by leaf as well as root inoculation in tomato seedlings. A transposon induced mutant has been isolated, which exhibit virulence deficiency through root inoculation in tomato seedlings but not by leaf inoculation. The linkage between the transposon insertion and the virulence phenotype has been proven using recreation of the mutation in the wild type by natural transformation. The whole genome sequencing has identified the transposon insertion in the *cyp450* gene homologue of F1C1 genome. Two independent mutations have been created in *cyp450* gene using different strategies and both mutants exhibit virulence deficiency by root inoculation in tomato seedlings. Both mutants exhibit a greater magnitude of virulence deficiency when inoculated by soil drenching but not by petiole pricking inoculation in grown up tomato plants. A full length *cyp450* gene amplified from the wild type genome complements the virulence deficiency in a mutant. The mutants exhibit a greater susceptibility than the wild type to tomato root extract. This corroborates the virulence deficiency exhibited by the *cyp450* mutants through root inoculation. Interestingly, the LC-MS study reveals the presence of several antibacterial compounds in the root extracts of tomato. To the best of our knowledge, this is the first demonstration of the role of *cyp450* in the virulence of plant pathogenic bacterium, *R. pseudosolanacearum*. Considering the known role of *cyp450* in metabolite degradation or biosynthesis of metabolites in organisms, its role in *R. pseudosolanacearum* virulence through root inoculation but not through shoot inoculation is interesting. This work also expands our understanding regarding the diversity of biological functions carried out by *cyp450* in bacteria. Cytochrome P450 (CYP) are heme-containing, divergent superfamily of monooxygenases distributed widely in organisms ranging from bacteria, plants, and humans (Permana et al. 2023). In plants, its copy number is found to be unusually high, where it is involved in various endogenous cellular processes, including xenobiotic degradation, environmental stress tolerance, secondary metabolite synthesis, and disease pathophysiology (Song et al. 2021). These enzymes may contribute to the virulence of plant-associated bacteria by facilitating the detoxification of plant defence compounds and altering metabolic pathways essential for survival within the plant (Coleman et al. 2011; Cresnar et al. 2011; Lah et al. 2011). Previous reports suggest involvement of *cyp450* genes in the virulence of many phytopathogenic fungi, like *Verticillium dahlia* and *Alternaria alternata* (Zhang et al. 2022; Fu et al. 2025). In these fungi, *cyp450* genes have been predicted to involve in detoxification of the oxygen free radicals inside the plant. The role of *cyp450* genes in the virulence of plant pathogenic bacteria has not been reported earlier. *R. pseudosolanacearum* F1C1 genome has a single copy of the *cyp450* gene present in a monocistronic operon. The differential virulence by the *cyp450* mutants in this study indicates its probable role in adapting to root tissue during infection. Earlier finding indicates that disease development in tomato seedlings by root inoculation is more complex than that inoculation in leaf (Kabyashree et al. 2021). It is known that root and shoot tissues are different in their physiology and metabolism (Zeiss et al. 2019). Plant tissues behave as distinct ecological niches for the pathogen. Before entering the xylem, pathogen encounters the rhizosphere and root cortex that are enriched with antimicrobial secondary metabolites, reactive oxygen species, phenolic compounds, flavonoids, etc. Once the pathogen moves inside the xylem, it has to transit into a comparatively nutrient limited but chemically different vascular environment. For successful pathogenesis, the pathogen requires dynamic metabolic adaptation towards different *in planta* environment. Successful pathogens remodel their metabolic pathways to exploit host nutrients while simultaneously detoxifying antimicrobial metabolites encountered during infection (Anderson 2023; Olive and Sassetti 2016). Root being associated with soil might be adapted to overcome the soil microorganisms by producing several antimicrobial compounds (Baetz and Martionia 2014), which the endophytic pathogen needs to overcome to successfully invade and colonize the root (Haicher et al. 2014). Cytochrome P450 might be helpful to phytopathogens and endophytes those invade through the root to overcome the hostile root environment by detoxifying the antimicrobials. The observation of greater sensitiveness of the *cyp450* mutants towards root extracts than the wild type F1C1 also indicate the possible adaptive role of *cyp450* in the root environment. The virulence deficiency in the mutants is more severe in the case of grown-up plants than in the seedlings. In the case of seedlings, the bacteria are directly exposed to the plant root surface, whereas in the case of grown-up plants, the bacteria are introduced in the soil surrounding the root through drenching. Therefore, the pathogen must overcome the soil metabolites as well as the root metabolites. In a preliminary study, we have observed that the root extract of grown-up plants contains greater antibacterial activity against *R. pseudosolanacearum* F1C1 than seedlings (unpublished data). Therefore, the severe phenotype might be attributed to the antibacterial activity of both the root, and the metabolites present in the soil, which can be studied in the future.

The methanolic root extract of tomato seedlings contains phenolic acid, flavonoids, polyphenols, terpenes, fatty acid-derived compounds, carboxylic acid, and amino acid derivatives, which has variety of roles in plant defence responses. A previous study suggests that these secondary metabolites act as defences against pathogenic microbes and animal pests. At elevated concentrations, many of these defence-related metabolites can inhibit or even kill microorganisms *in vitro* (Wang et al. 2020; Yang et al. 2021). Malic acid present in the root extract has been reported to inhibit the growth of *R. solanacearum* FJAT-91 (Chen et al. 2025). In a similar context, gallic, coumaric, caffeic, hydroxybenzoic, chlorogenic, and ferulic acids were reported to have growth suppression activity against *R. solanacearum* (Hamed and Messiha 2018; Li et al. 2021). Previously it has been demonstrated that hydroxycinnamic acids (HCAs), a plant-derived phenolic compound, is present in the tomato plant (Leonard et. al 2017; Lowe power et. al 2015; Zeiss et. al 2019). We have also observed the presence of HCA in the root extract of tomato seedlings. *R. solanacearum* mutants having disruption of the *fcs* gene, encoding feruloyl-CoA synthetase required for the degradation of HCA is reduced in virulence on tomato plants by inoculation through the root. The *fcs* mutant was found to have increased sensitivity to the caffeate and p-coumarate as well as reduced early colonization of tomato roots. This study indicates that HCA degradation is particularly important during the early stages of infection, when the pathogen encounters plant-derived phenolic compounds in the rhizosphere and roots (Lowe power et. al 2015). Studies have also suggested that cytochrome P450 systems have been associated with detoxification of plant defence compounds (Vitas et al. 1999). Collectively, these studies indicated that the ability of *R. solanacearum* to metabolize plant-derived antimicrobial compounds is required for successful colonization and infection. In our study, we have demonstrated the inhibitory effect of tomato root extracts, whereas the leaf extracts did not exhibit any inhibitory effect towards any of the strains. Notably, the *cyp450* mutants exhibited greater sensitivity to tomato root extracts in comparison to the wild-type F1C1 strain. LC-MS analysis further revealed the presence of several antibacterial compounds in the tomato root extract. These findings suggest that the *cyp450* gene may contribute to the ability of *R. pseudosolanacearum* F1C1 to tolerate the hostile root environment and play a role during virulence through root of tomato plants. To our knowledge, this is the first study to demonstrate differential sensitivity of *R pseudosolanacearum* mutants to tomato root extracts compared with the wild-type strain. It will be interesting to find out the growth inhibitory effect of any specific metabolites in the root extract towards mutant and wild type. Screening different tomato cultivars with the mutant and a comparative study of their resistant in future will be helpful regarding the usefulness of this study in finding out bacterial wilt resistance. Further study will explore the mechanistic role of *cyp450* in other phytopathogens or endophytes that invade the host through the root. Phytopathogens like *Xanthomonas*, which infect through the leaf, it will also be interesting to understand the role of *cyp450* in their virulence.

The function of cytochrome P450 in rhizobia suggest that they may play specialized roles during plant–microbe interactions. In *Rhizobium* sp. BR816, a *cyp450* gene was found to be highly expressed in bacteroid within root nodules and was induced under microaerobic and acidic conditions that resemble the nodule environment, suggesting a role in adaptation to the plant host rather than in general bacterial growth or nitrogen fixation (Luyten et al. 2001). This finding further indicates that bacterial *cyp450* may also contribute to plant-associated adaptations. Earlier studies have demonstrated that the virulence phenotype of *R. solanacearum* can vary depending on the mode of inoculation. Direct inoculation into the plant xylem through petiole inoculation has previously been used to investigate virulence in *R. solanacearum* (Khokhani et al. 2018). Mutations in two genes involved in the flagellar biosynthesis pathway resulted in reduced virulence following root inoculation but not after shoot inoculation, indicating that these genes are required during the early stages of host invasion but are dispensable once the pathogen reaches the xylem (Tans-Kersten et al. 2001). Similarly, mutations in two genes *cheA* and *cheW* involved in the chemotaxis signal transduction pathway caused reduced virulence after root inoculation but not after shoot inoculation, further supporting that chemotaxis is important for root invasion but is not required after vascular colonization (Yao and Allen 2006). Mutation in the *pilT* gene, encoding an ATPase required for type IV pilus dynamics shows significant reduction in virulence through root inoculation but not through shoot inoculation (Singh et al. 2018b). In all previously reported cases, the gene products are surface-associated bacterial proteins. However, in the present study, the involvement of CYP450, which seems to be an intracellular protein, their role in differential virulence is intriguing. *R. solanacearum* genome has many adhesins, such as hemagglutinins, which have been predicted to have a role in the host range of this pathogen (Genin and Denny 2012). It will be interesting to find out if any of those adhesins exhibit differential virulence in host plant tissues.

## 4 Experimental Procedures

### 4.1 Bacterial strains, plasmids, and growth conditions

The bacterial strains used in this study and their characteristics are listed in Table 1. Plasmids used and created for this study are listed in Table 2. *R. pseudosolanacearum* F1C1 (Kumar et al. 2013) was grown at 28°C in BG (Bacterial Growth) medium (Plener et al. 2010) and minimal medium (Ray et al. 2015). BG medium consists of 1% peptone, 0.1% casamino acid, 0.1% yeast extract, supplemented with 0.5% glucose. Triphenyl tetrazolium chloride (0.05%) and 1.5% agar were added for BG agar plates. The minimal medium composition is as follows in g/L: 1.25×10^−4^ FeSO_4_.7H_2_O; 0.5 (NH_4_)_2_SO_4_; 0.05 MgSO_4_.7H_2_O; 3.4 KH_2_PO_4_. KOH was used to adjust the pH 7.0. *E. coli* strains were grown at 37°C using LB medium and LB supplemented with 1.5% agar medium (Ausubel et al. 1989). To grow *R. pseudosolanacearum* F1C1 and *E. coli* the growth medium was supplemented with antibiotics whenever needed, with the following concentrations: Ampicillin (Amp; 50 μg/ml), Spectinomycin (Spec; 50 μg/ml), Gentamycin (Gen; 50 μg/ml), Kanamycin (Kan; 50 μg/ml). The chemicals, antibiotics, and media components used in the study were purchased from Hi-Media, Mumbai, India, and SRL, New Mumbai, India. Glasswares and plastic wares were purchased from Borosil and Tarsons, Kolkata, India.

**TABLE 1.**
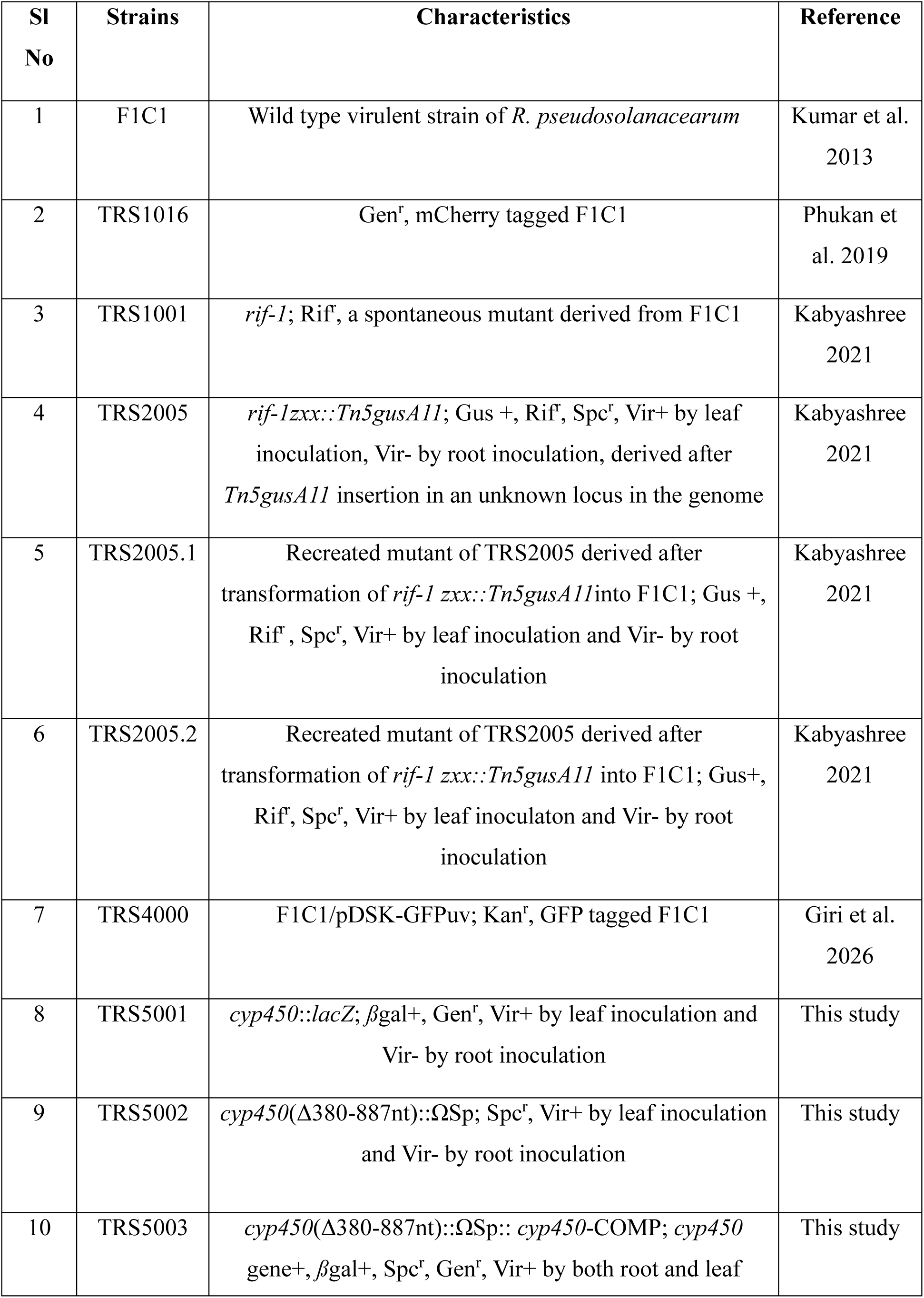
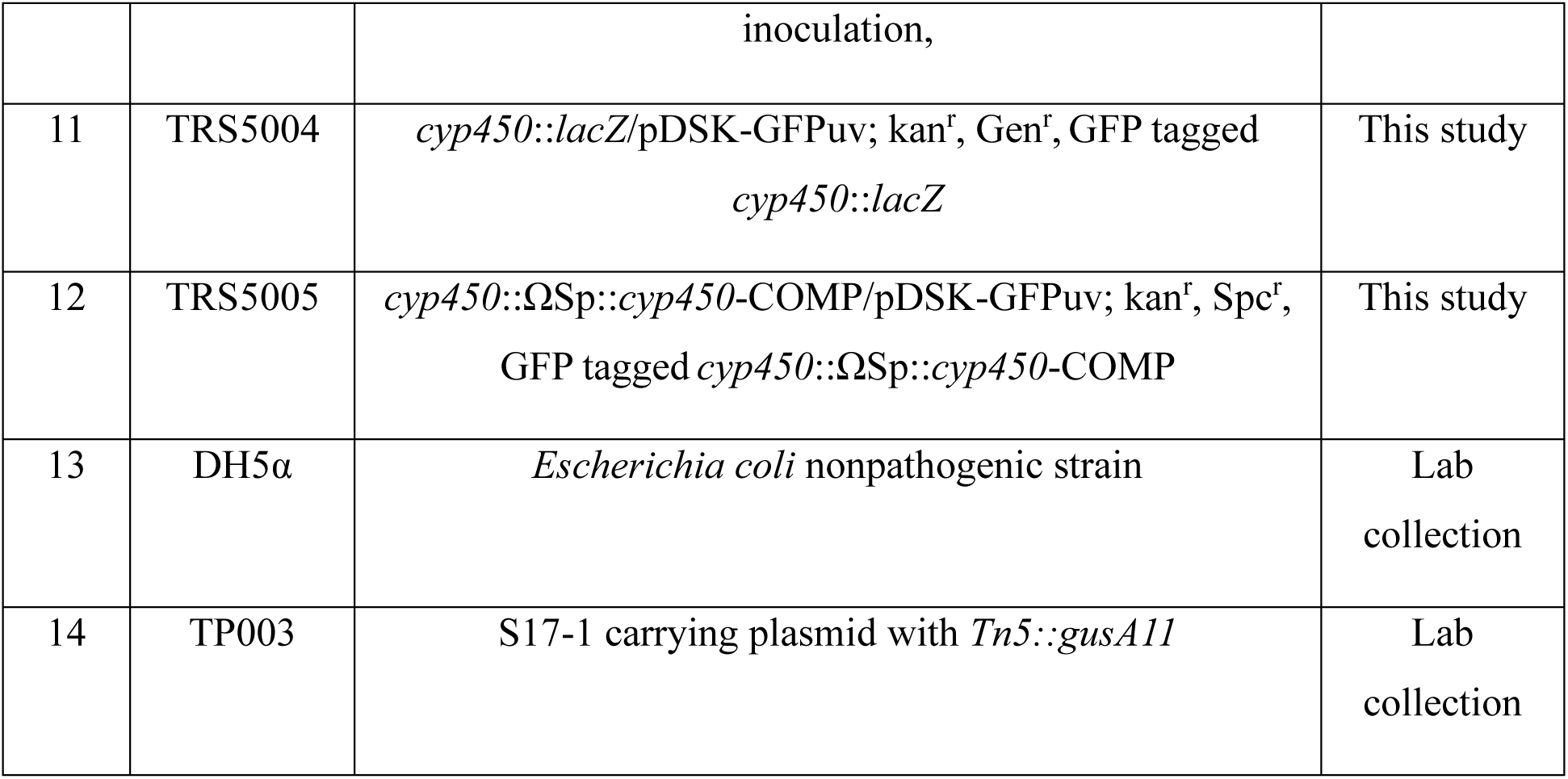
List of bacterial strains used in this study.

**TABLE 2.** List of plasmids used in this study.

| SI No | Plasmids | Characteristics | Reference |
| --- | --- | --- | --- |
| 1 | pGEMT | Amp<sup>r</sup>, TA Cloning vector | Promega |
| 2 | pCZ367 | Amp<sup>r</sup>, Gen<sup>r</sup>, Insertional vector with *lacZ* reporter | Cunnac et al. 2004 |
| 3 | PHP45 | Spc<sup>r</sup>, plasmid carrying a 2 kb Ω spectinomycin cassette | Prentki 1984 |
| 4 | pLD001 | [pGEMT::*cyp450*(250-938nt)]; pGEMT carrying 688 bp<br>*cyp450* gene fragment, Amp<sup>r</sup> | This study |
| 5 | pLD002 | [pCZ367::*cyp450*(250-938nt)]; pCZ367 carrying 688 bp<br>*cyp450* gene fragment, Amp<sup>r</sup>, Gen<sup>r</sup> | This study |
| 6 | pLD003 | [pGEMT::*cyp450* (*cyp450* -269 to -1 nt upstream and +1<br>to +379 nt of 5' *cyp450* gene)]; pGEMT carrying 648 bp<br>amplified region from F1C1 genome, Amp<sup>r</sup> | This study |
| 7 | pLD004 | [pGEMT::*cyp450* (886 -1197 nt of 3' *cyp450* gene-immediate downstream to *cyp450* 1 - 438 nt)]; pGEMT<br>carrying 749 bp amplified region from F1C1 genome,<br>Amp<sup>r</sup> | This study |
| 8 | pLD005 | [pGEMT::*cyp450* (*cyp450* -269 to -1 nt upstream and +1 to +379 nt of 5' *cyp450* gene)- *cyp450* (886 -1197 nt of 3' *cyp450* gene-immediate downstream to *cyp450* 1 - 438 nt)]; pGEMT carrying 648 bp (fragment I) and 749 bp (fragment II) amplified region from F1C1 genome, Amp<sup>r</sup> | This study |
| 9 | pLD006 | [pGEMT::*cyp450* (*cyp450* -269 to -1 nt upstream and +1 to +379 nt of 5' *cyp450* gene)-ΩSp-*cyp450* (886 -1197 nt of 3' *cyp450* gene-immediate downstream to *cyp450* 1 - 438 nt)]; pGEMT carrying 648 bp (fragment I), 2 kb Ω Spectinomycin cassette and 749 (fragment II) bp amplified region from F1C1 genome, Amp<sup>r</sup>, Spc<sup>r</sup> | This study |
| 10 | pLD007 | (pCZ367::*cyp450*-complementaion); pCZ367 carrying 1346 bp region that contains entire 1197 bp *cyp450* gene, 88 bp upstream (-88 to -1 nt) covering the promoter region and 61 bp immediate down stream of *cyp450*, Amp<sup>r</sup>, Gen<sup>r</sup> | This study |

### 4.2 Restriction enzymes, ligation and polymerase chain reaction

The ligase, restriction enzymes, and all other reagents used throughout this study were used as per the manufacturer’s recommendations. A volume of 20 µl of PCR reaction was set up as following reagents: 10X dream taq Buffer 2.0 µl, dNTPs 1.2 µl (2mM), DMSO 0.9 µl, each oligo 1.0 µl (10 pmol), template DNA (5-10 ng), 1.0 µl; dream-Taq DNA polymerase 0.2 μl (5 U/µl), final volume of 20 µl was adjusted with molecular grade water. The ligation reactions were performed in 1:3 (vector: insert) ratio for overnight at 4°C. The buffers, restriction enzymes, Pfu polymerase, DNA polymerase, DNA ligases and other components used in this study were purchased from Thermo Fisher Scientific, Waltham, MA, USA.

### 4.3 *cyp450*::*lacZ* insertion mutation

A schematic representation of the strategy for insertion mutation is provided in Figure S2a. *R. pseudosolanacearum* F1C1 contains one *cyp450* gene of size 1197 bp. A part of it, 688 bp within (250 - 938 nt) was amplified from the genomic region using the oligo oLD0709F1: GGGAAGCTTTGCTCAATCCGCCCATGCAC and oLD0709R1: GGGTCTAGAAGCATCAGGAACACCGTCGT. *HindIII* and *XbaI* restriction sites were already included at the 5́ end of F1 and R1 oligos, respectively. The amplified fragment was gel purified (Qiagen, Hilden, Germany) and cloned into pGEMT (Promega, Madison, Wisconsin, United States) to obtain the plasmid pLD001. The transformants were selected on ampicillin containing LB agar medium. The transformant was initially confirmed by colony PCR and recombinant plasmid pLD001 was isolated by using plasmid isolation kit (Qiagen, Hilden, Germany). Nested PCR using oligos oLD0709F1: GGGAAGCTTTGCTCAATCCGCCCATGCAC and oLD0709R3 GGGTCTAGACGTTCGGCATCCGCGACGGC (Table 3) was performed to confirm the amplified fragment cloned was from the desired region. The cloned 688 bp was digested out from the plasmid pLD001 using *HindIII* and *XbaI* and was cloned into pCZ367 plasmid vector (Cunnac et al. 2004, Plener et al. 2010, Ray et al. 2015) at the *HidIII* and *XbaI* site. This is a reporter gene fusion vector where the reporter gene *lacZ* is located downstream of the inserted fragment. After successful cloning in pCZ367, the plasmid pLD002 was confirmed by colony PCR and restriction digestion with *HindIII* and *XbaI*. The constructed plasmid pLD002, was transformed into wild type F1C1 by heat shock mediated transformation method (Cowell et al. 2026, Giri et al. 2026) and the transformants were selected on gentamycin containing BG medium. In the transformants it is expected the entire plasmid construct has been integrated by single homologous recombination as pCZ367 cannot replicate inside *Rps* F1C1. pLD002 integration in the genome will result into the disruption of the gene *cyp450*. To confirm that the integration has been in *cyp450* locus and not in any random location of the genome, a PCR was performed using an oligo of *lacZ* gene Lac_R: GGGTGGCGAAAGGGGGATGTGCT and another oligo from the upstream of *cyp450* gene. Lac_F2: GGGTCCTGGCTTTATGAAACTCG (Table 3) to obtain a successful amplification of 1287 bp. The *cyp450* insertion mutant *cyp450*::*lacZ* (TRS5001), was considered for virulence study as well as sensitiveness towards the tomato root extract.

**TABLE 3.**
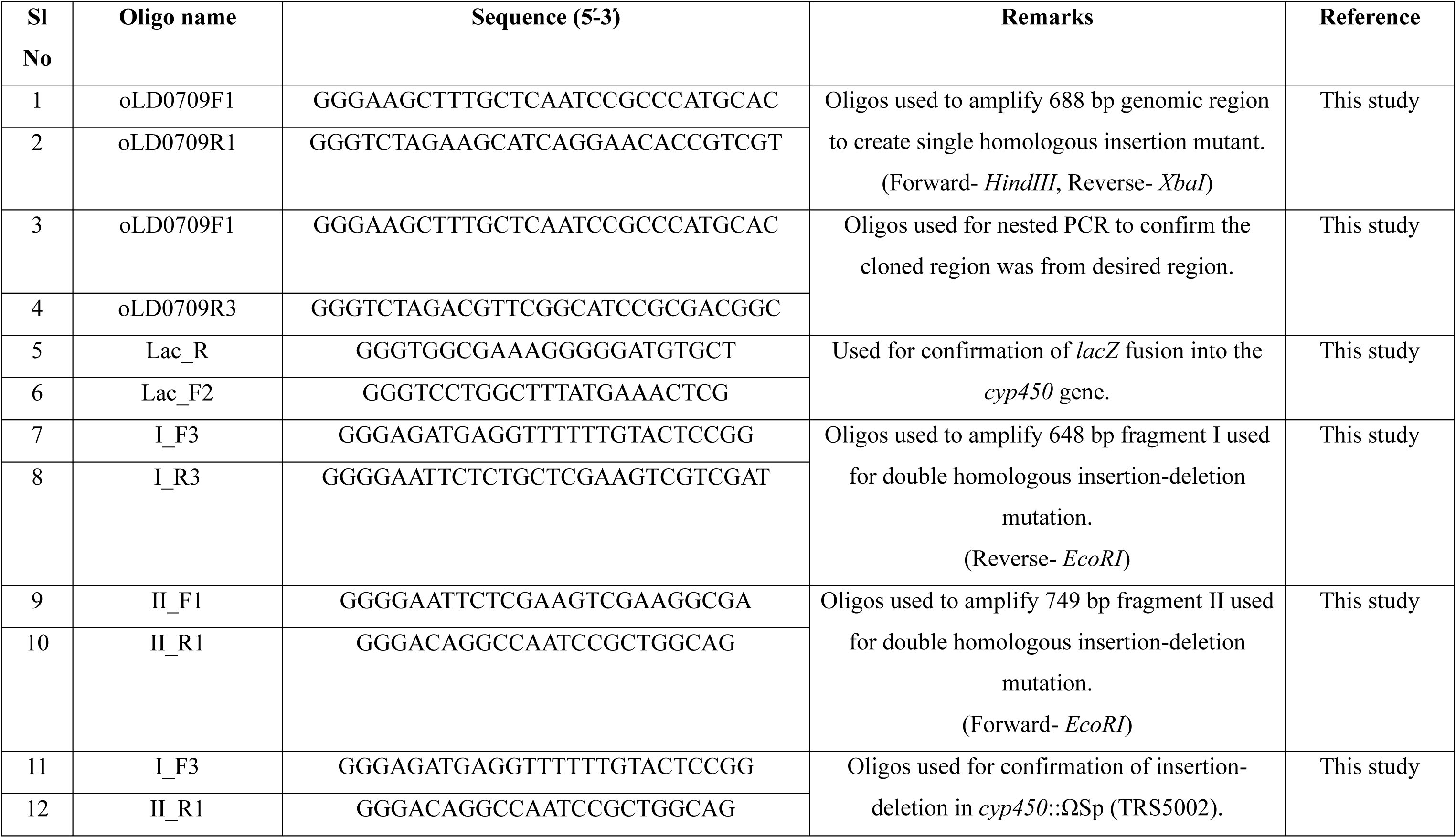
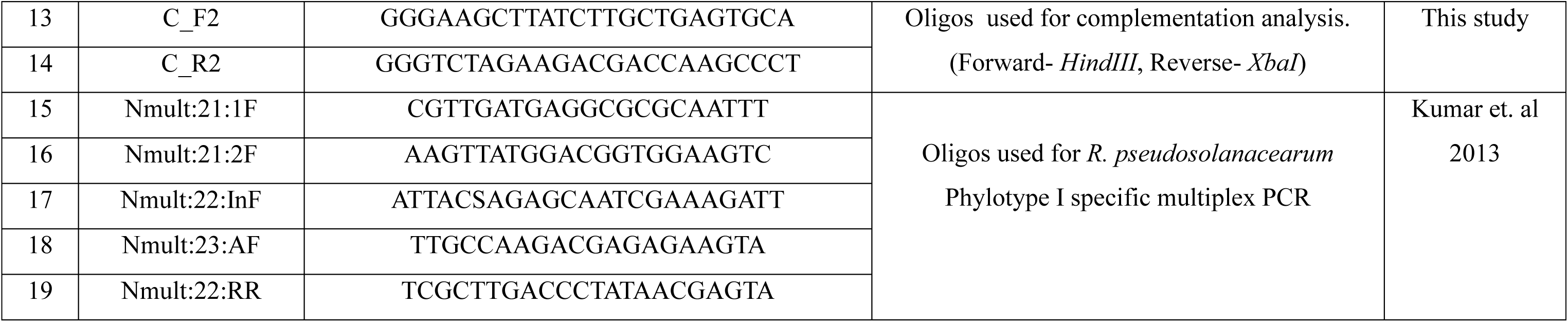
List of oligos used in this study.

**TABLE 4.**
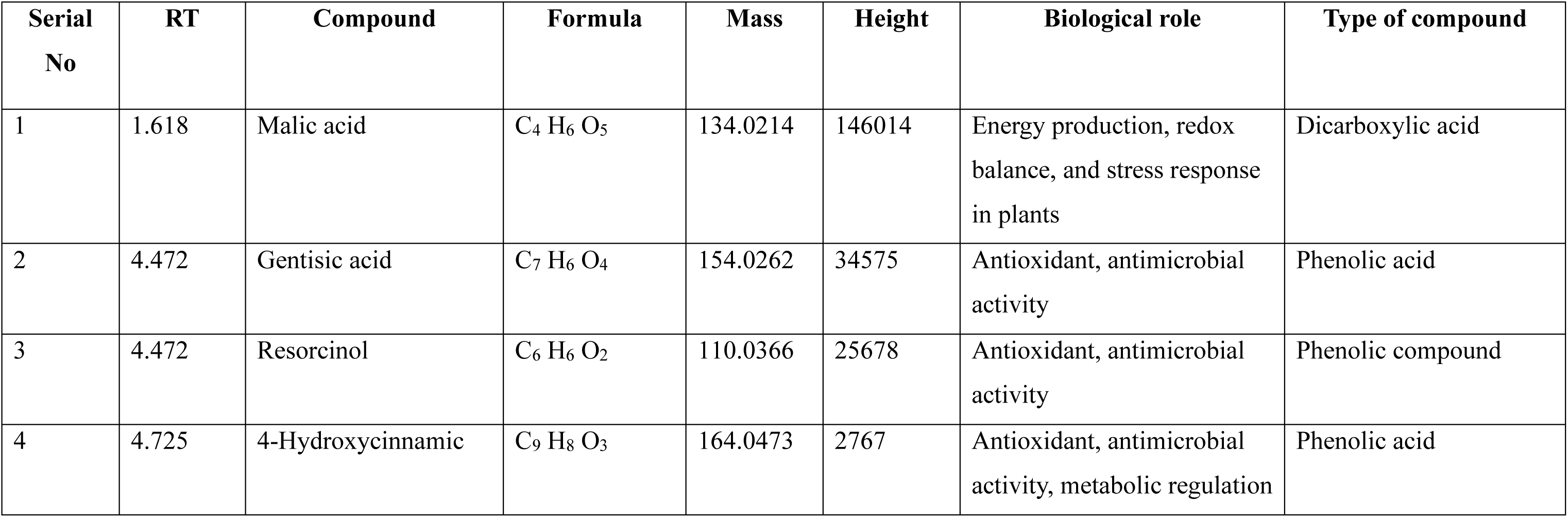
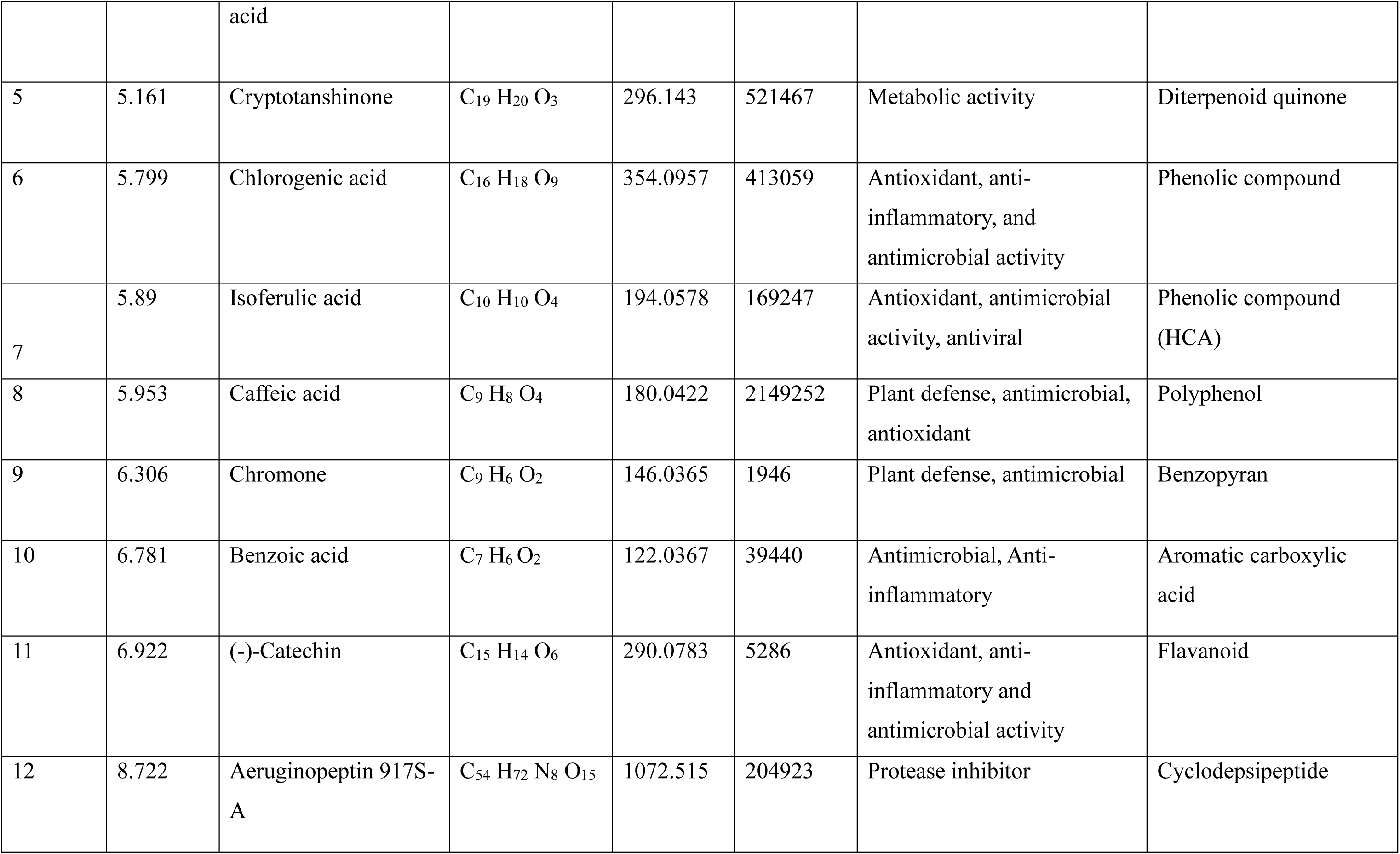
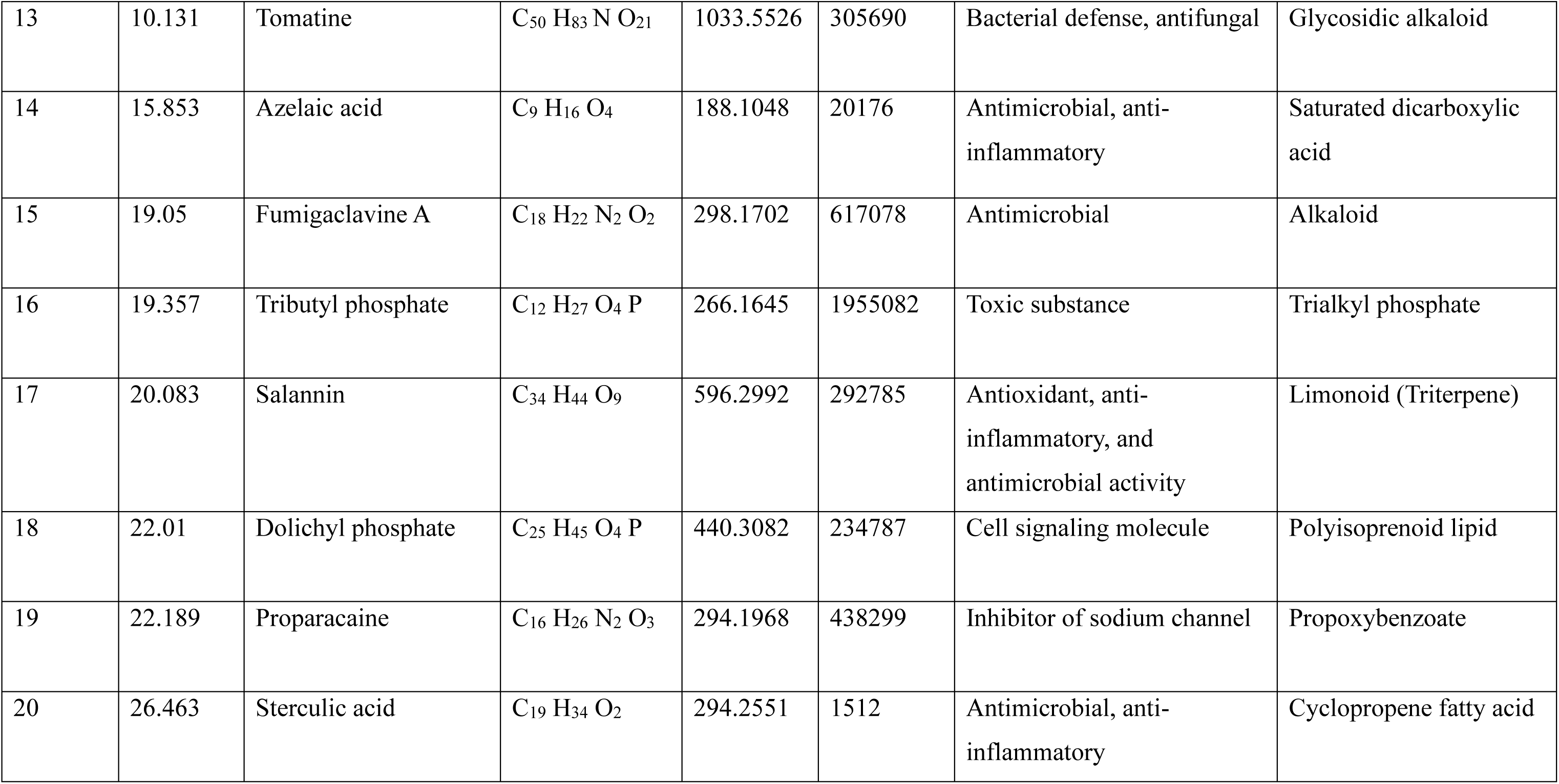
Antibacterial compounds identified in methanolic root extracts of tomato seedlings by HR-LCMS analysis.

All the oligos used in this study were obtained from BIOLINKK (New Delhi, India) and the list of oligos are mentioned in Table 3.

### 4.4 *cyp450*::ΩSp insertion deletion mutation

A schematic representation of the insertion-deletion strategy is provided in Figure S2b. Fragment I of size 648 bp, which covers -269 to -1 bp upstream regions of the *cyp450* gene and +1 to +379 bp 5́ region of *cyp450* gene, was amplified by using oligos I_F3:GGGAGATGAGGTTTTTTGTACTCCGG and I_R3:GGGGAATTCTCTGCTCGAAGTCGTCGAT. An *EcoRI* restriction site was incorporated at the 5́ end of the reverse oligo (I_R3) to facilitate subsequent cloning. Fragment II of size 749 bp, comprising 311 bp (886 to 1197 nt) from the 3́ region of the *cyp450* gene and 438 bp immediate downstream regions of the gene was amplified by using oligos II_F1:GGGGAATTCTCGAAGTCGAAGGCGA and II_R1:GGGACAGGCCAATCCGCTGGCAG. An *EcoRI* restriction site was incorporated at the 5́ end of the forward oligo (II_F1) to facilitate subsequent cloning. Fragment I and fragment II was cloned separately into pGEMT vector to obtain recombinant plasmids pLD003 and pLD004, respectively. Cloning was confirmed by colony PCR and *EcoRI* restriction digestion. pLD003 was also digested with *EcoRI* and *NcoI* to confirm the orientation of the cloning for fragment I in pGEMT. pLD004 was digested with *EcoRI* and *NdeI* to confirm the orientation of the cloning for fragment II. Fragment II was isolated from pLD004 after restriction digestion with *EcoRI* and *NdeI*, and was cloned with pLD003 which was also restriction digested with *EcoRI* and *NdeI* to create pLD005 (total 4397 bp), where fragment I (379 bp) and fragment II (311 bp) joined in such a way that it created a 507 bp gap in the *cyp450* gene. The Ω cassette was excised from plasmid pHP45 (Prentki and Krish 1984; Ray et al 2015) by digesting with *EcoRI*, and was ligated with pLD005 digested with *EcoRI*, to create pLD006. The complete construct was, 3 kb pGEMT vector-648 bp fragment I-2kb Ω spectinomycin cassette -749 bp fragment II (total 6397 bp). The recombinant plasmid pLD006 was linearized by *NdeI* digestion and introduced into F1C1 by natural transformation (described below). Transformants were selected on spectinomycin containing medium, and spectinomycin resistant colony arising from the double homologous recombination were screened. The insertion-deletion mutation in the *cyp450* gene was confirmed by a PCR amplification using oligos I_F3: GGGAGATGAGGTTTTTTGTACTCCGG and II_R1: GGGACAGGCCAATCCGCTGGCAG (Table 3). The oligos amplify the entire *cyp450* gene together with 269 bp of the upstream region and 438 bp of the downstream flanking region yielding an expected amplification of 1904 bp (∼ 2 kb) from the wild type F1C1 genome but not from the transformant genome. The *cyp450* insertion-deletion strain *cyp450*::ΩSp (TRS5002), was considered for virulence study as well as sensitiveness towards the tomato root extract.

### 4.5 Complementation Analysis

A schematic representation of the strategy for complementation analysis is provided in Figure S3. The fragment 1346 bp which encompasses the entire 1197 bp of *cyp450* open reading frame, the upstream 88 bp (-88 to -1 nt) comprising its probable promoter region and 61 bp immediate downstream of the *cyp450* gene was amplified using the error free pfu polymerase. The oligos used were C_F2: GGGAAGCTTATCTTGCTGAGTGCA and C_R2: GGGTCTAGAAGACGACCAAGCCCT having *HindIII* at the 5́ end of F2 and *XbaI* at the 5́ end of R2 respectively. The amplified fragment was double digested with *HindIII* and *XbaI* and ligated with pCZ367 which was also double digested with *HindIII* and *XbaI*. This ligated plasmid created recombinant plasmid pLD007. The plasmid pLD007 was introduced by heat shock transformation into the virulence deficient insertion-deletion strain *cyp450*::ΩSp (TRS5002). The transformant was selected on gentamycin and spectinomycin supplemented BG plates. The transformant was named as complemented strains TRS5003 (*cyp450*::ΩSp::*cyp450*-COMP). PCR amplification was performed using oligo C_F2: GGGAAGCTTATCTTGCTGAGTGCA and C_R2: GGGTCTAGAAGACGACCAAGCCCT, where an amplification was found in *cyp450*::ΩSp::*cyp450*-COMP (TRS5003) but not in *cyp450*::ΩSp (TRS5002). It indicated a full length *cyp450* restored in the genome of *cyp450*::ΩSp::*cyp450*-COMP (TRS5003). *cyp450*::ΩSp::*cyp450*-COMP (TRS5003) was studied for *ß*-Galactosidase production, virulence by soil drenching, sensitiveness towards root extract.

### 4.6 *R. pseudosolanacearum* F1C1 natural transformation method

Natural transformation in *R. pseudosolanacearum* F1C1 was performed following the method described in Ray et al. (2015) and Phukan 2018. *Rps* F1C1 was grown for overnight at 28°C in BG medium supplemented with glucose. A volume of 1 ml from the overnight grown culture was taken and centrifuged for 10 min at 4000 rpm at room temperature. The supernatant was discarded and the pellet was resuspended in 1.0 ml sterile distilled water. Thereafter, 100 ul of the suspension (1% of the culture) was added to 10,0 ml of the medium supplemented with 15% glycerol as a sole carbon source. The culture was incubated for 48 h at 28°C. The growth of the bacterium was visible from the increase in the turbidity. 15 μl of the F1C1 culture was mixed with 4 μl (∼2 μg) of linearized plasmid pLD006 and then spotted on the sterile nitrocellulose membrane placed on the BG-agar plates (without glucose). The plates were incubated for 48 h at 28°C. The grown bacterial cells on the nitrocellulose membrane were scraped using toothpick and resuspended in 150-200 μl of sterile distilled water. The suspension was spread on spectinomycin-supplemented BG agar plates. After incubating for 48 h at 28°C, the plates were observed for transformed colonies.

### 4.7 *lacZ* reporter gene expression study

The *β*-Galactosidase activity was performed according to Miller (1972) and Ray et al. (2015). The experiment was carried out in two different growth media, i.e., BG and minimal medium. The mutant *cyp450*::*lacZ* (TRS5001), the complemented strain *cyp450*::ΩSp::*cyp450*-COMP (TRS5003) and wild-type F1C1 cells were grown separately in BG and minimal medium overnight at 28°C. 2.0 ml of culture from each sample was centrifuged at 4°C at 13000 rpm for 2 min. The pellet was resuspended with 750 μl of Z buffer, and the final volume was made 1.5 ml. The OD was measured at 600 nm, considering the Z buffer as blank. 750 μl of the bacterial suspension was taken and to it, 100 μl chloroform and 50 μl 0.1% SDS were added. The suspension was mixed by vertexing and incubated for 10 minutes at 28°C. Thereafter, 150 μl of ONPG (ortho-Nitrophenyl-β-D-Galactopyranoside) was added in each tube and incubated at 37° C. The time of ONPG addition was recorded. Once solution turned yellow, the reaction was terminated by adding 375 μl of 1M Na_2_CO_3_ solution and incubated in ice. The tubes were then centrifuged at 13000 rpm to settle down the debris and absorbance of the supernatant was measured at 420 nm and 550 nm. The *β*-Galactosidase enzyme activity was measured in miller units for both BG and minimal medium (Miller 1972).

### 4.8 Virulence study in tomato seedlings

Virulence study (*Solanum lycopersicum* Pusa Ruby, Durga Seeds, India) was performed by leaf inoculation (Kumar et al. 2017) as well as root inoculation (Singh et al. 2018) in seven-day-old tomato seedlings. Bacterial inoculum with concentration ∼ 10^9^ CFU/ml was prepared following the methods described in Kabyashree et al. (2020). Three independent experiment was performed. In each experiment two sets consisting of 40 seedlings were inoculated with wild-type F1C1, *cyp450*::*lacZ* (TRS5001), *cyp450*::ΩSp (TRS5002), *cyp450*:: ΩSp::*cyp450*-COMP (TRS5003), and water as control. The inoculated seedlings were kept in a growth chamber (Orbitech, India), maintaining a temperature of 28°C, relative humidity at 75%, and a photoperiod of 12 h (day/night). Scoring was recorded up to 10 days post-inoculation (dpi) for disease progression (Kabyashree et al 2020). The scoring was done based on the number of seedlings wilted each day. The disease scoring was given as: for the appearance of symptoms, score 1, blackening of the stem to wilting of both the cotyledon leaves, score 2, complete wilting of the seedling, score 3. The percentage of wilting was calculated, and survival probability was calculated using Kaplan-Meier Survival Probability Test (Kaplan et al. 1958), and for this analysis, the wilted seedling was scored as 1, whereas score 0 represents the healthy seedlings.

### 4.9 Virulence study in grown-up tomato plants by soil-drenching as well as petiole pricking

To study the virulence of *cyp450* mutants through root of tomato plants, the soil drenching method of bacterial inoculation was performed as described in Giri et al. (2025). Tomato plants were grown for one month in the greenhouse at 28°C, 12h photoperiod (day/night). Two independent experiments were performed. In each experiment, 20 grown up plants per strain such as wild-type F1C1, mutants *cyp450*::*lacZ* (TRS5001) and *cyp450*::ΩSp (TRS5002), complemented strain *cyp450*::ΩSp::*cyp450*-COMP (TRS5003) and sterile distilled water as control were used. Bacterial suspensions of all the strains were prepared and OD_600_ were adjusted to 0.6. A volume of 50 ml of the respective bacterial inoculum per plant was poured into the soil in each pot. For the control group, 50 ml of sterile distilled water was poured into the soil. Plants were maintained in the greenhouse (28°C, 12h photoperiod day/night) and their growth was monitored regularly until 20 dpi. Disease scoring was recorded daily from 0 for no disease to 4 for complete wilting (Ray et al. 2015). To validate Koch’s postulates, the pathogen was re-isolated from the F1C1 and *cyp450*::ΩSp::*cyp450*-COMP (TRS5003) inoculated wilted plants by the ooze streaming test and confirmed by PCR amplification by the *cyp450* gene specific oligos in both strains (Abo-Elyousr et al. 2024).

To study the virulence of *cyp450* mutants through shoot inoculation in tomato plants, the petiole pricking method of inoculation was performed. Like the above, two independent experiment was performed. In each experiment 12 plants per strain such as wild-type F1C1, mutants *cyp450*::*lacZ* (TRS5001) and *cyp450*::ΩSp (TRS5002), complemented strain *cyp450*::ΩSp::*cyp450*-COMP (TRS5003) and sterile distilled water as control was used. In each plant all leaf petioles except those of the top leaves were inoculated by pricking. A sterile sewing machine needle was used for inoculation in the petiole of the plant (Diatloff et al. 2000). The sterile needle was dipped in a bacterial culture having ∼10^9^ CFU/ml of bacterial concentration and each petiole was pricked twice. In the case of control plants, the needle was dipped in sterile distilled water. The inoculated plants were monitored regularly for disease symptom development until 15 dpi and disease progression was assessed by recording the disease score. Usually there were three to five compound leaves with as many petioles in each plant, which were inoculated that many times (Ingel et al. 2022). Each compound leaf inoculated at the petiole was observed for the wilting symptoms. Scoring was done for each leaf and the average score was used for each plant. The wilting of the entire plant was observed faster in case of petiole pricking than in comparison to soil drenching.

### 4.10 Sensitivity of Rps F1C1 towards tomato root and leaf extracts

The seven days old cotyledon stage seedlings were used for this assay. The roots of the seedlings were cut near the root-shoot junction and 0.5 g of roots were measured for further analysis. The roots were surface sterilized with 0.04% bavistin and 0.04% mercuric chloride solution, followed by 70% ethanol, as described in Kabyashree et al. (2020). Stringent surface sterilization was used to avoid any bacterial contamination. An aliquot of water from the final rinse was plated on BG agar plate to observe no growth of microorganisms to confirm proper surface sterilization process and to use as a control in this assay. The roots were crushed in a sterile mortar-pastel using sterile distilled water (500 μl) until a uniform homogenized solution appeared. The homogenate was transferred into a sterile microcentrifuge tube and further studied its antibacterial properties against wild-type F1C1, mutants *cyp450*::*lacZ* (TRS5001), *cyp450*::ΩSp (TRS5002) and *cyp450*::ΩSp::*cyp450*-COMP (TRS5003). Three wells of 4.0 mm diameter each were made in the BG-agar plate using a sterile cork borer, with one well positioned at the center of the plate and the other two wells located 2 cm away from the central well. Wild type F1C1 and mutants were streaked close to the well at an equal distance of 1 cm above and below. Followed by it, 50 µl of the root extract was added to the middle well, while 50 µl of the sterile distilled water was added which was taken from the distilled water washing step of surface sterilization. The plates were incubated at 28°C for 48 h. The experiment was repeated for three times. For the agar well diffusion method, 100 µl of bacterial cultures (∼10^9^ CFU/ml) was spread on a BG-agar plate. A well of 4.0 mm diameter well was made with a sterile cork borer at the center of the BG-agar plate to which 50 µl of root extract was added. The plate was incubated at 28°C for 48 h. Observation of a clear zone surrounding the well indicated the antagonistic activity of the root extract. The zone of clearance was measured, and the area was also calculated for each strain. The experiment was repeated for three times for each strain.

Similarly, the leaf extract sensitivity assay of tomato seedlings was performed. Tomato leaves were cut using a scissor and 0.5 g of leaf was collected for further analysis. The leaf samples were surface sterilized and homogenized in sterile distilled water following the protocol described above. The sensitivity of wild type F1C1, *cyp450* mutants and complemented strain was assessed using the streaking method as described above. The experiment was repeated three times. As no inhibition was observed by the leaf extract towards any strain, so the well diffusion method was not performed.

### 4.11 Metabolomics study of tomato root extracts using HR-LCMS/MS

An untargeted metabolomic approach was used to identify metabolites present in the root extract of tomato seedlings. The methanolic extract of root samples was dried and subjected to HR-LCMS/MS (MS Q-TOF G6550A) analysis to identify antibacterial compounds present in it. For chromatographic separation Hypersil GOLD C18 (2.1 x 100 mm 3-micron) column was used with a solvent gradient 0.1% formic acid in water (A) and 100% acetonitrile (B), flowing at 0.300 ml/min. The gradient started with B at 5% at 1 min, increasing to 100% at 25 min, remaining at 100% until 30 min, then returning to 5% at 31 min and held until 35 min, with a total run time of 35 min. The QTOF conditions were set to Auto MS/MS mode, with specific parameters including Dual AJS ESI ionization mode, a drying gas temperature of 250°C, drying gas flow of 13 l/min, nebulizer pressure at 35 psig, vaporizer/sheath gas temperature of 300 °C, and sheath gas flow at 11 l/min. The Vcap was set to 3500 V, fragmentor to 175 V, skimmer to 65 V, and Octopole RF Peak 750 V. The precursor per cycle maximum was 10, with an absolute threshold of 10000 counts and a relative threshold of 0.01%. The mass spectra were acquired over an m/z range of 120–1200.

### 4.12 Presence of bacteria in tomato seedlings

Presence of bacteria in seedlings after root or leaf inoculations was studied by crushing the surface sterilized inoculated seedlings and by performing a dilution plating in BG medium containing antibiotics. Seven-day old tomato seedlings were inoculated separately with wild type F1C1, *cyp450*::*lacZ* (TRS5001), *cyp450*::ΩSp (TRS5002) and complemented strain *cyp450*::ΩSp::*cyp450*-COMP (TRS5003). For both the root and leaf inoculation bacterial inoculum with concentration 10^9^ was used. At 4 dpi from each group of infected seedlings, one seedling was selected and surface sterilized by dipping in 70% ethanol for 1 min followed by washing twice with sterile distilled water for 1-2 min (Phukan, 2019). After that seedling was transferred into a microcentrifuge tube and homogenized separately using sterile micro pestle (Abdos, India). The homogenate was resuspended in 1 ml of sterile distilled water from which it was serially diluted till 10^7^ dilutions. From each dilution five 10 µl aliquots were spotted onto BG agar plates supplemented with appropriate antibiotics. The plates were incubated at 28°C for 48 h, after which the colonies observed were counted.

Mixed inoculation experiment was performed in three different combinations such as mCherry F1C1 (TRS1016) + *cyp450*::ΩSp (TRS5002), mCherry F1C1 (TRS1016) + *cyp450*::ΩSp::*cyp450*-COMP (TRS5003), *cyp450*::ΩSp (TRS5002) + *cyp450*::ΩSp::*cyp450*-COMP (TRS5003). For selection purpose mCherry-marked *R. pseudosolanacearum* (TRS1016) strain was used which was gentamycin resistant (Monteiro et al. 2012; Capela et al. 2017; Phukan et al. 2019). The saturated culture (∼10^9^ CFU/ml) of the two strains were mixed with equal volume and inoculated in tomato seedlings by root inoculation. Bacterial population was quantified at 0 dpi, 2 dpi and 4 dpi in the seedlings after surface sterilization. On 0 dpi, two seedlings were sampled, on 2 dpi another two seedlings were sampled and on 4 dpi another two seedlings were sampled. The selected seedlings were then surface sterilized and transferred into a sterile microcentrifuge tube. The seedlings were homogenized in a sterile micro pestle and mixed with 1 ml of sterile distilled water. The homogenate was serially diluted up to 10^7^ dilutions and from each dilution five 10 µl aliquot was spotted in BG agar plates supplemented with appropriate antibiotics. The plates were incubated in 28°C for 48 h, after which colonies were counted and CFU/ml was calculated. The same procedure was followed for all treatments and CFU/ml, ratio of both the strain was calculated.

Bacterial presence in inoculated seedlings was further performed by visualization green fluorescence protein (GFP) marked bacterial strains. The GFP-plasmid pDSK-GFPuv, which carries the GFP and a kanamycin resistance gene (Wang et al. 2007). The GFP tagged in the wild type F1C1 (TRS4000) was used in this study (Giri et al. 2026). The mutant *cyp450*::*lacZ* (TRS5001) and complemented strain *cyp450*::ΩSp::*cyp450*-COMP (TRS5003) were tagged with plasmid pDSK-GFPuv. Seven-day old tomato seedlings were inoculated separately with the GFP tagged strains by root inoculation. Seedlings inoculated with sterile distilled water and those inoculated with non-GFP tagged strain were included as negative control. At 4 dpi the roots, stems and leaves of inoculated seedlings were harvested, surface-sterilized, and examined under a fluorescence microscope (EVOS FL, Life Technologies) to visualize colonization.

### 4.13 Motility assays

Twitching motility was performed using a glucose-free BG agar plate following the method described by Bhuyan et al. (2024). The bacterial culture of F1C1, *cyp450*::*lacZ* (TRS5001), *cyp450*::ΩSp (TRS5002) *cyp450*::ΩSp::*cyp450*-COMP (TRS5003) was grown overnight in BG medium. The culture was serially diluted to 10^4^-fold, and a volume of 5 μl was spotted on the glucose-free BG agar plates. After incubating at 28°C, the plates were observed under an inverted fluorescence microscope with a 4X objective lens (40X magnification) within a 12-18 h duration of time.

Swimming motility was performed in a semisolid glucose-free BG agar (0.2% agar) plate as described by Ray et al. 2015. To visualize the growth of the bacteria, 0.05% tetrazolium chloride (TZC) was added to the medium as a growth indicator. A sterile toothpick was dipped in the bacterial culture and carefully touched in the middle of the soft agar plates and incubated at room temperature. After 12 h of incubation, swimming motility was observed by the formation of a circular halo zone.

### 4.14 Biochemical characterization

The mutants, complemented strains along with the wild type F1C1 was screened for extracellular enzyme production such as cellulase followed by the method described in Barman et al. (2017) and Kumar 2014.. For cellulase enzyme production 80 μl of bacterial culture (∼ 10^9^ CFU/ml) was added into the well prepared in the middle of carboxymethyl cellulose medium supplemented with 1.5% agar plates. The plates were incubated at 28°C for 48 h. After incubation, the plates were washed with distilled water to remove the bacterial growth around the well and 1% Congo red solution was added for 48 h and incubated at room temperature. After that, the plates were destained with 1 M NaCl and kept overnight at room temperature to observe a distinct clear zone for positive cellulase production.

The mutants, complemented strains along with the wild type F1C1 was screened for pectinase production followed by the method described in Giri et al. (2025). For the pectinase assay, freshly grown bacterial culture (∼10^9^ CFU/ml) was added to the well prepared in the middle of the BG agar plates supplemented with 0.5% pectin and incubated at 28°C for 3 days. After incubation, 2% cetyltrimethylammonium bromide solution was added for 30 min, and then the plates were washed with 0.5 M NaCl to visualize the clear zone around the well for positive pectinase production. The generation time of TRS5001, TRS5002, TRS5003 and wild type F1C1 was calculated following the method described in Bhuyan et al. (2023). The oxidative stress response of the mutants was studied against different concentrations of H_2_O_2_ as described in Zhang et al. (2022).

The hypersensitive response of the mutants was studied by following the method described in Amaria et al. (2023). The Nicotiana rustica variety of tobacco was used and 100 µl, saturated bacterial cell suspension (∼10^9^ CFU/ml) was infiltrated into the lower surface of the tobacco leaves using a sterile syringe. The inoculated leaves were observed after 24 to 48 h to visualize the necrotic lesions in the infiltrated regions.

### 4.15 Data Analysis

The generation of graphs and statistical analyses were performed using the software GraphPad Prism and Origin.

## Acknowledgements

We are extremely grateful for the anonymous reviewers whose critical observation and insightful comments greatly improved the quality, clarity and overall presentation of this manuscript. We sincerely thank Prof. Stéphane Genin from INRAE, Toulouse, France for providing the pCZ367 and pHP45 plasmid vector. We sincerely thank Prof. Kirankumar S. Mysore from Oklahoma State University, US for providing the pDSK-GFPuv plasmid vector. We are grateful to Prof. Marc Valls for the mCherry plasmid vector. We thank Dr Mohit Yadav for his valuable suggestions. We acknowledge DBT, GoI UExcel NER grant to SKR. We further acknowledge the PhD thesis work of Dr Kristi Kabyashree (http://hdl.handle.net/10603/336047). We are thankful to Priya Rani Daimari and Zoomani Rabha for their comments on the manuscript. LD expresses gratitude to DST, GoI, New Delhi, for the INSPIRE-JRF/SRF fellowship. SJG and SB1 are thankful for the JRF fellowship from the DBT, GoI, New Delhi, under grant (BT/PR41637/NER/95/1753/2021) awarded to SKR. SK1 acknowledges the JRF fellowship from the UGC-CSIR, GoI, New Delhi. MJ acknowledges the JRF fellowship from the UGC-CSIR, GoI, New Delhi. SB2 is thankful for fellowship from the DBT, GOI, New Delhi, granted under (BT/PR54976/BSA/33/428/2024) awarded to SKR SKR, SK2 and AK are thankful to the ANRF-PAIR grant. SKR expresses gratitude to Prof. R. V. Sonti for the transposon mutagenesis and Dr. S. Genin for the targeted insertion mutagenesis lessons. We are thankful to all the pervious Scholars in the SKR laboratory who set up the seedlings inoculation experiments. The authors were thankful to Sophisticated Analytical Instrumental Facility (SAIF), IIT Bombay, for HR-LCMS analysis of the root extract.

## Author Contribution

**LD** performed the experiments, analyzed the data, prepared the draft manuscript, reviewed, and edited. **KK** performed the experiment, screening and identification of *cyp450* gene, reviewed, and edited. **SJG** performed experiment, data analysis, reviewed, and edited the article**. KB** performed genome sequence analysis, reviewed, and edited the article. **DC, SB1, SK1, MJ and SB2** performed formal data analysis, reviewed the manuscript. **SK2, AK and GJ** critical analysis and reviewed the manuscript. **SKR,** and **PBP** supervised, reviewed, edited, and conceptualization.

## Data Availability Statement

All the data generated during this study are included in the manuscript and supplementary files.

