## Supplemental Files for "Mutation in *Ralstonia pseudosolanacearum* F1C1 cytochrome *p450* homologue impairs virulence through root but not through shoot of tomato plants"

^4^ Present address: Indian Institute for Human Settlements, Bengaluru, 560080, India

**TABLE S1. Studying the presence of *cyp450* mutants in seedlings by performing mix inoculation with the wild type F1C1**

| **0 dpi** | | | | | |
| --- | --- | --- | --- | --- | --- |
| **Inoculation** | **Sample** | **Set 1 CFU/ml** | **Ratio** | **Set 2 CFU/ml** | **Ratio** |
| Wild type+Mutant | mCherryF1C1 | 1.4 × 10^3^ | 0.7 | 6 × 10^3^ | 4.3 |
|  | *cyp450*::ΩSp | 2 × 10^3^ |  | 1.4 × 10^3^ |  |
| Wild type + Complemented strain | mCherryF1C1 | 4 × 10^2^ | 2 | 2 × 10^2^ | 1 |
|  | *cyp450*::ΩSp::*cyp450*-COMP | 2 × 10^2^ |  | 2 × 10^2^ |  |
| Mutant+ Complemented strain | *cyp450*::ΩSp | 8 × 10^4^ | 4 | 4 × 10^4^ | 2 |
|  | *cyp450*::ΩSp::*cyp450*-COMP | 2 × 10^4^ |  | 2 × 10^4^ |  |
| **2 dpi** | | | | | |
| **Inoculation** | **Sample** | **Set 1 CFU/ml** | **Ratio** | **Set 2 CFU/ml** | **Ratio** |
| Wild type+Mutant | mCherryF1C1 | 1 × 10^8^ | 0.25 | 1.2 × 10^8^ | 2 |
|  | *cyp450*::ΩSp | 4 × 10^8^ |  | 6 × 10^7^ |  |
| Wild type + Complemented strain | mCherryF1C1 | 6 × 10^6^ | 3 | 1 × 10⁶ | 2.5 |
|  | *cyp450*::ΩSp::*cyp450*-COMP | 2 × 10^6^ |  | 4 × 10⁵ |  |
| Mutant+ Complemented strain | *cyp450*::ΩSp | 8 × 10⁵ | 10 | 6 × 10⁵ | 5 |
|  | *cyp450*::ΩSp::*cyp450*-COMP | 8 × 10^4^ |  | 1.2 × 10⁵ |  |
| **4 dpi** | | | | | |
| **Inoculation** | **Sample** | **Set 1 CFU/ml** | **Ratio** | **Set 2 CFU/ml** | **Ratio** |
| Wild type+Mutant | mCherryF1C1 | 2 × 10⁵ | 0.1 | 4 × 10⁶ | 0.5 |
|  | *cyp450*::ΩSp | 2 × 10⁶ |  | 8 × 10⁶ |  |
| Wild type + Complemented strain | mCherryF1C1 | 4 × 10⁷ | 1 | 2 × 10⁶ | 10 |
|  | *cyp450*::ΩSp::*cyp450*-COMP | 4 × 10⁷ |  | 2 × 10⁵ |  |
| Mutant+ Complemented strain | *cyp450*::ΩSp | 2 × 10⁵ | 0.28 | 2 × 10⁵ | 0.25 |
|  | *cyp450*::ΩSp::*cyp450*-COMP | 7 × 10⁵ |  | 8 × 10⁵ |  |

**TABLE S2. Studying the presence of bacteria in an inoculated seedling**

| **Root inoculation (4 dpi)** | |
| --- | --- |
| **Samples** | **CFU/ml** |
| F1C1 | 2.6 × 10^8^ |
| *cyp450*::*lacZ* (TRS5001) | 8 × 10⁶ |
| *cyp450*::ΩSp (TRS5002) | 2.2 × 10^6^ |
| *cyp450*::ΩSp::*cyp450*-COMP (TRS5003) | 4 × 10^4^ |
| **Leaf inoculation (4 dpi)** | |
| **Samples** | **CFU/ml** |
| F1C1 | 8 × 10^6^ |
| *cyp450*::*lacZ* (TRS5001) | 2 × 10^5^ |
| *cyp450*::ΩSp (TRS5002) | 1.4 × 10^5^ |
| *cyp450*::ΩSp::*cyp450*-COMP (TRS5003) | 2 × 10^7^ |

**Figure Legends**

**FIGURE S1 | Virulence phenotype of transposon induced mutant TRS2005 and recreated mutants of TRS2005**

Transposon induced mutant TRS2005 and the recreated mutants of TRS2005 (TRS2005.1 & TRS2005.2) along with wild type F1C1 were inoculated in tomato seedlings by root and leaf inoculations. In case of leaf inoculation, the pathogenicity magnitudes of mutant as well as the recreated mutants were similar to that of F1C1. In case of root inoculation, both these mutant (TRS2005) and recreated mutants (TRS2005.1 & TRS2005.2) were significantly reduced in virulence than F1C1. The experiments were performed in triplicates and every time, three sets of 30 seedlings each were inoculated.

**FIGURE S2 | Strategy used for insertion mutation *cyp450*::*lacZ* and insertion-deletion mutation *cyp450*::ΩSp**

Schematic representation of homologous recombination based *cyp450* gene targeting strategies that demonstrates (a) creation of insertion mutant *cyp450*::*lacZ* (TRS5001) via single homologous recombination and (b) creation of insertion–deletion mutant *cyp450*::ΩSp (TRS5002) via double homologous recombination.

**FIGURE S3 | Strategy used for complementation analysis**

Schematic representation of complementation analysis used for creation of complemented strain *cyp450*::ΩSp::*cyp450*-COMP (TRS5003)

**FIGURE S4 | Virulence phenotype of mutants in tomato seedlings**

40 seedlings per strain were inoculated through root and leaf. Symptoms were monitored and scored till 10 dpi. (a) The X-axis represents days post inoculation (dpi), and the Y-axis represents the percentage of survival probability. Survival probability of tomato seedlings inoculated with *cyp450*::*lacZ* (TRS5001) is 75.67% (p=0.0086), survival probability of tomato seedlings inoculated with *cyp450*::ΩSp (TRS5002) is 67.56% (p=0.0405), survival probability of tomato seedlings inoculated with *cyp450*::ΩSp::*cyp450*-COMP (TRS5003) is 47.64% (p=0.6766), and seedlings inoculated with F1C1 were 45.94%. (b) For leaf inoculation, after 10 dpi, the survival probability of tomato seedlings inoculated with *cyp450*::*lacZ* (TRS5001), *cyp450*::ΩSp (TRS5002), and *cyp450*::ΩSp::*cyp450*-COMP (TRS5003) was observed to be 18.53% (p=0.7342), 10.13 % (p=0.8680), 4.76 % (p=0.4151) respectively and seedlings inoculated with F1C1 were 11.26 %. *cyp450* mutants *cyp450*::*lacZ* (TRS5001) and *cyp450*::ΩSp (TRS5002) are virulence deficient by root inoculation but not by leaf inoculation.

**FIGURE S5 | Virulence phenotype of *cyp450* mutants in grown-up tomato plants by soil drenching**

In the second independent experiment of soil drenching, Figure (a), 20 plants per strain were inoculated by soil drenching method. 50 ml of the respective bacterial inoculum (OD_600_=0.6) per plant was poured into the soil in each pot. The photograph was taken at 20 dpi. 14 plants wilted in case of wild type F1C1, 14 plants wilted in case of complemented strain *cyp450*::ΩSp::*cyp450*-COMP (TRS5003). 6 plants wilted in case of *cyp450*::*lacZ* (TRS5001) and 1 plant wilted in case of *cyp450*::ΩSp (TRS5002) representing significant reduction in virulence compared to wild type and complemented strain. The all 20 plants in the control were healthy. For easy visualization of the disease, the pots were arranged properly in case A, The random occurrence of the disease as well as the escapees among the inoculated plants are schematically presented in figure (b) as shown by filled circle, healthy plants in green and wilted plants in brown. The disease progression is represented in Figure (c). Plants were monitored regularly and wilting symptoms were scored till 20 dpi at a disease index scale from 0-4. The graph represents one independent experiment consisting 20 plants per strain.

**FIGURE S6 |** **Virulence phenotype of *cyp450* mutants in grown-up tomato plants by petiole pricking**

In the second independent experiment of petiole pricking, (a) 12 plants per strain such as wild-type F1C1, *cyp450*::*lacZ* (TRS5001), *cyp450*::ΩSp (TRS5002), *cyp450*::ΩSp::*cyp450*-COMP (TRS5003) (10^9^ CFU/ml) and water control were inoculated by petiole pricking method. The photograph was taken at 15 dpi. All the 12 plants inoculated with the F1C1 and complemented strain developed wilting. 11 plants wilted in case of *cyp450*::*lacZ* (TRS5001) and 11 plant wilted in case of *cyp450*::ΩSp (TRS5002). Plants inoculated with sterile distilled water remained healthy throughout the experiment. The disease development is schematically presented by the color filled circles as greens represents healthy plants, brown represents wilted plants and yellow represents plants having wilting symptoms in Figure (b). The disease progression is represented disease in Figure (c).

**FIGURE S7 | Confirmation of the presence of wild type F1C1 and complemented strain in wilted plants by ooze steaming test.**

Presence of *R. pseudosolanacearum* F1C1 and *cyp450*::ΩSp::*cyp450*-COMP (TRS5003) was confirmed by spreading the collected ooze on triphenyl tetrazolium chloride (TTC) supplemented BG agar plate and *cyp450* gene-specific PCR amplification.

**FIGURE S8 | Antibacterial activity of tomato seedlings root extracts**

Study of bacterial growth regarding inhibition of tomato root extracts. In BG agar medium wild type F1C1, *cyp450* mutants and complemented strain (~10^9^ CFU/ml) were spread separately. One well of 4.0 mm diameter, positioned at the center of the plate was made. 50 µl of the tomato root extract was added to the middle well. The zone of inhibition was measured after 48 h of incubation at 28^o^ C. (a) Control plate, 50 µl of sterile distilled water was tested to check any inhibition activity (b) Zone of inhibition for wild-type F1C1, 24.6 ± 2.5. (c) Zone of inhibition for mutant *cyp450*::*lacZ* (TRS5001), 29.3 ± 1.5 (p=0.0209). (d) Zone of inhibition for mutant *cyp450*::ΩSp (TRS5002), 29 ± 1 (p=0.0320) (e) Zone of inhibition for complemented strain *cyp450*::ΩSp::*cyp450*-COMP (TRS5003), 23 ± 1 (p=0.6546) (f) Zone of inhibition for *E. coli* taken as negative control, 11.5 ± 0.7 (p= <0.0001). (g) the graph shows the statistical significance of zone of inhibition for all the tested strains. Tomato seedlings root extracts show greater inhibitory activity against *cyp450* mutants.

**FIGURE S9 | Study for bacterial presence in inoculated seedlings using GFP-tagged strains**

GFP-tagged strains such as mutant *cyp450*::*lacZ*/pDSK-GFPuv (TRS5004), complemented strain *cyp450*::ΩSp::*cyp450*-COMP/pDSK-GFPuv (TRS5005) and wild type F1C1/pDSK-GFPuv (TRS4000) was inoculated in tomato seedlings through root. Seedlings inoculated with non-GFP tagged strain *cyp450*::*lacZ* (TRS5001) and seedlings inoculated with sterile distilled water was taken as control. At 4 dpi, the fluorescence micrograph was captured using a fluorescence microscope (Life Technologies EVOS FL) with 100X magnification and a scale bar of 400 μm. Presence of wild type, mutant and complemented strains could be observed by visualizing the fluorescence in GFP tagged strains but not in the control seedlings. *cyp450* mutants are present inside tomato seedlings inoculated through root.

**FIGURE S10 | Extracellular enzymes and motility study of *cyp450* mutants, complemented strain and wild type F1C1**

The figure shows qualitative extracellular enzyme production such as pectinase and cellulase, motility study such as swimming and twitching motility in wild type F1C1, *cyp450*::*lacZ* (TRS5001), *cyp450*::ΩSp (TRS5002) and complemented strain *cyp450*::ΩSp::*cyp450*-COMP (TRS5003).

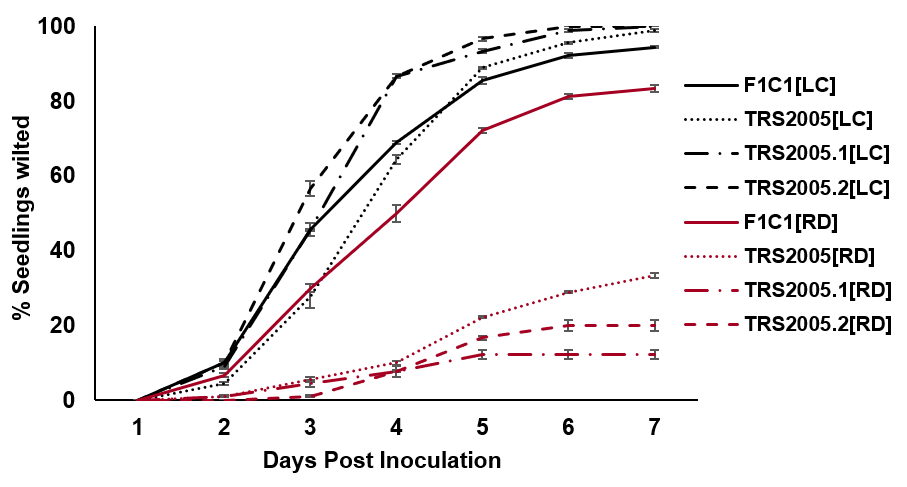

**Figure S1**

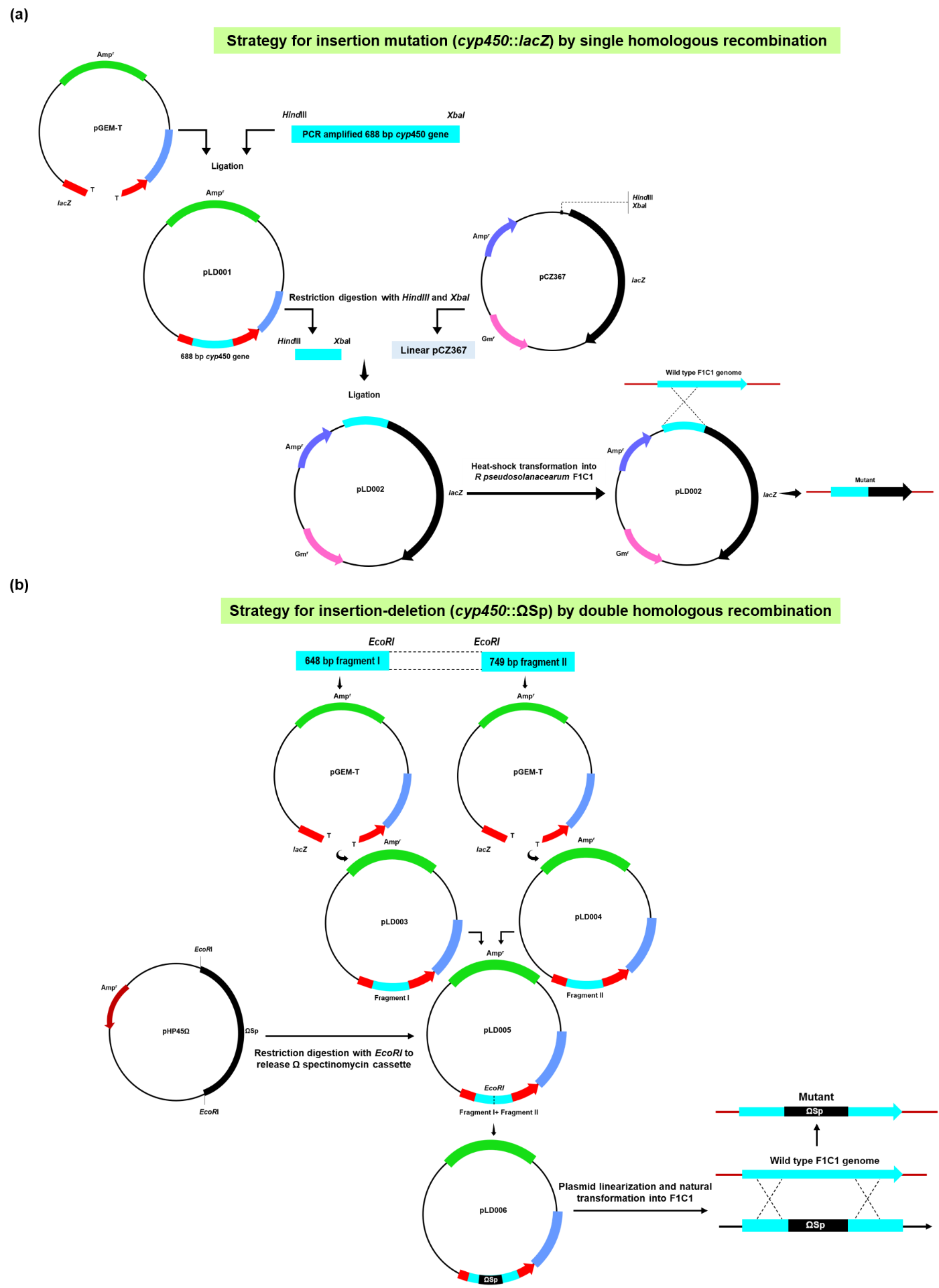

**Figure S2**

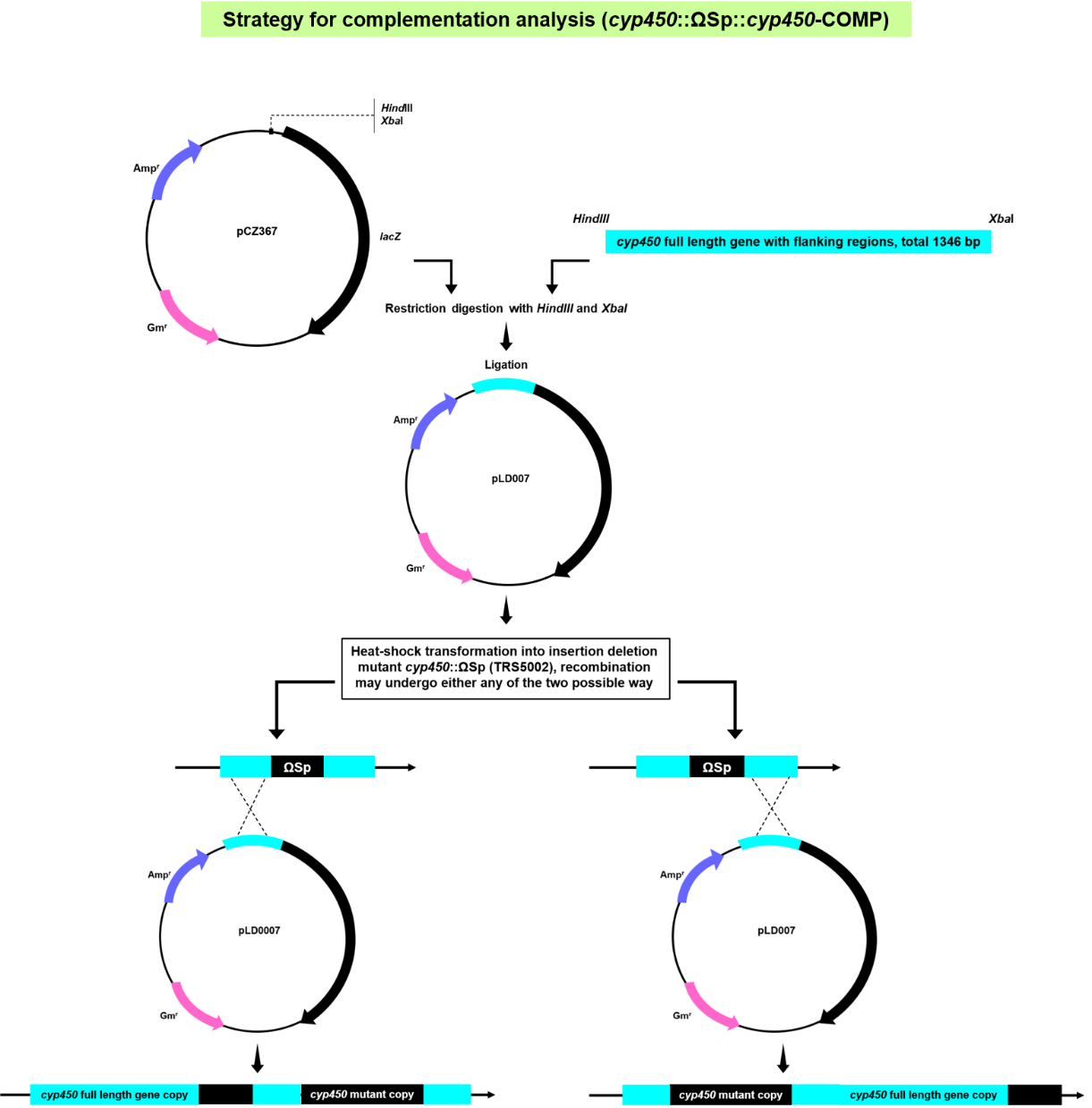

**Figure S3**

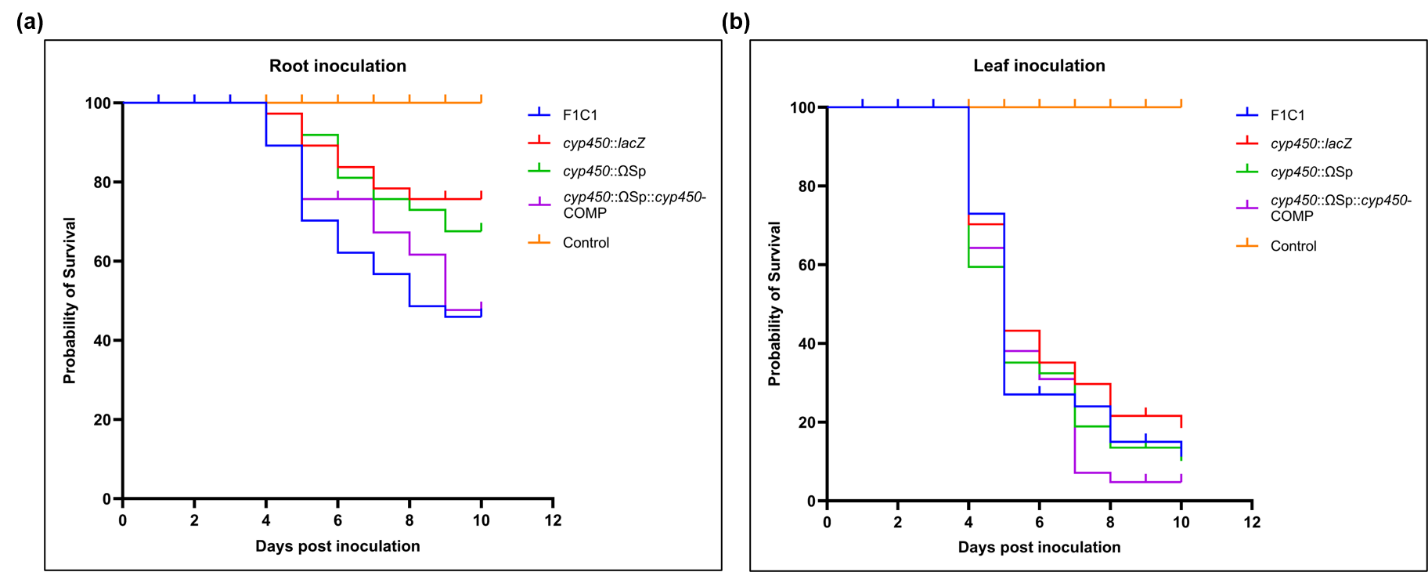

**Figure S4**

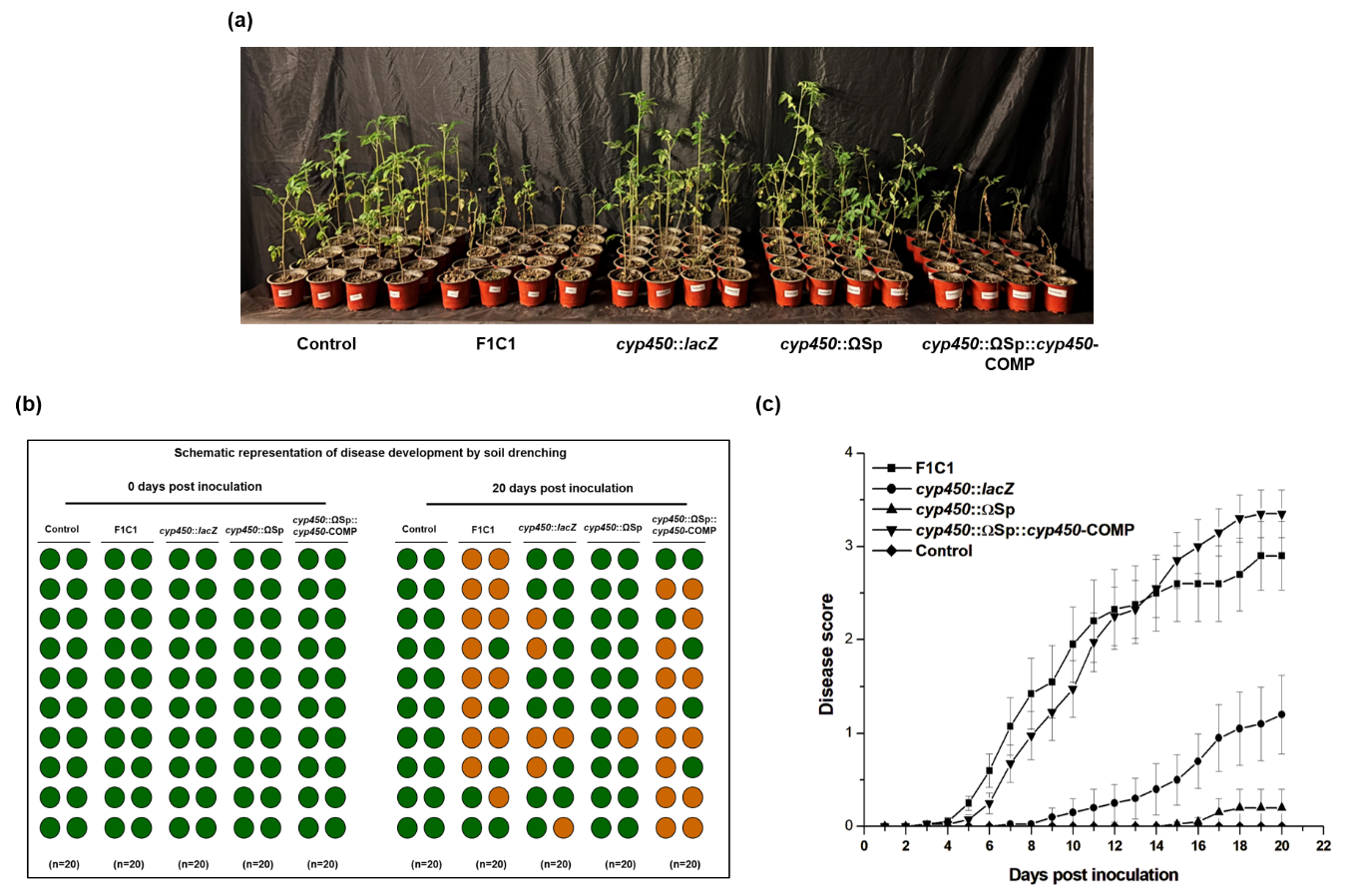

**Figure S5**

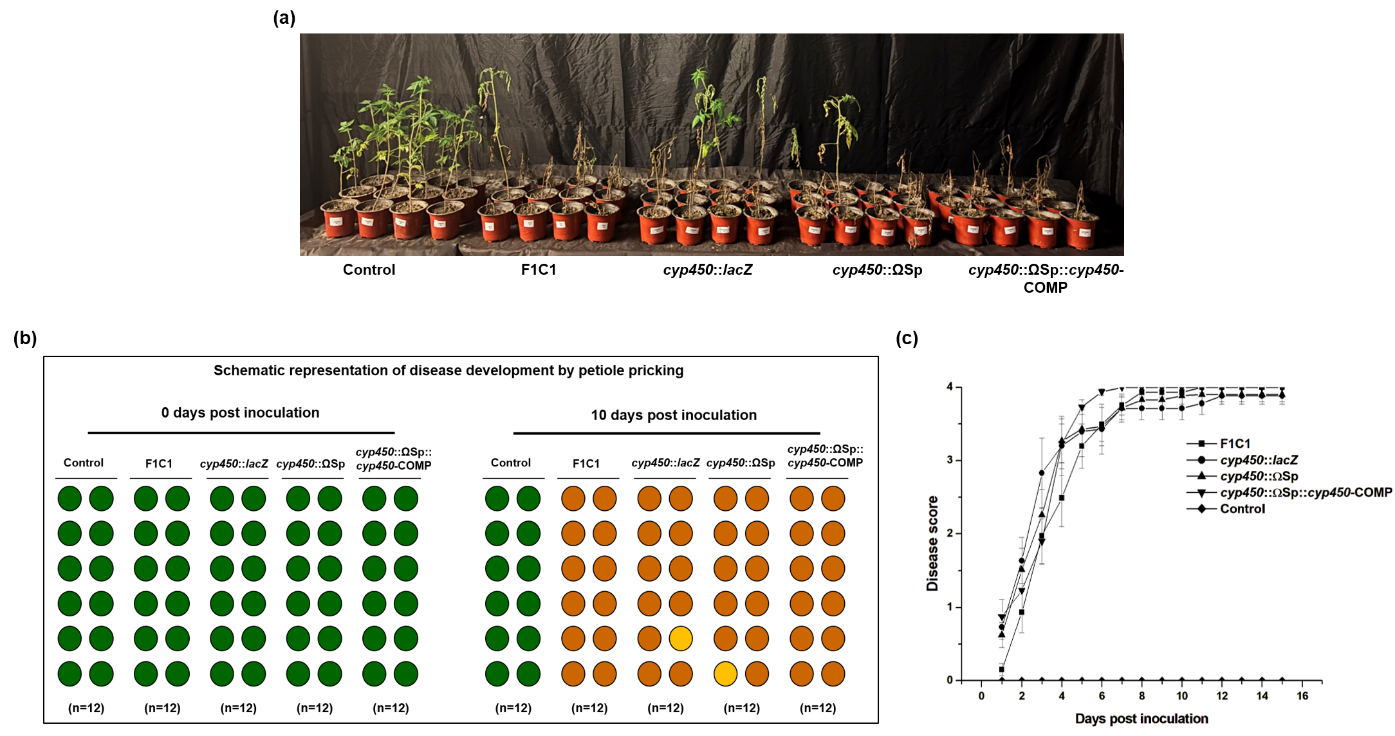

**Figure S6**

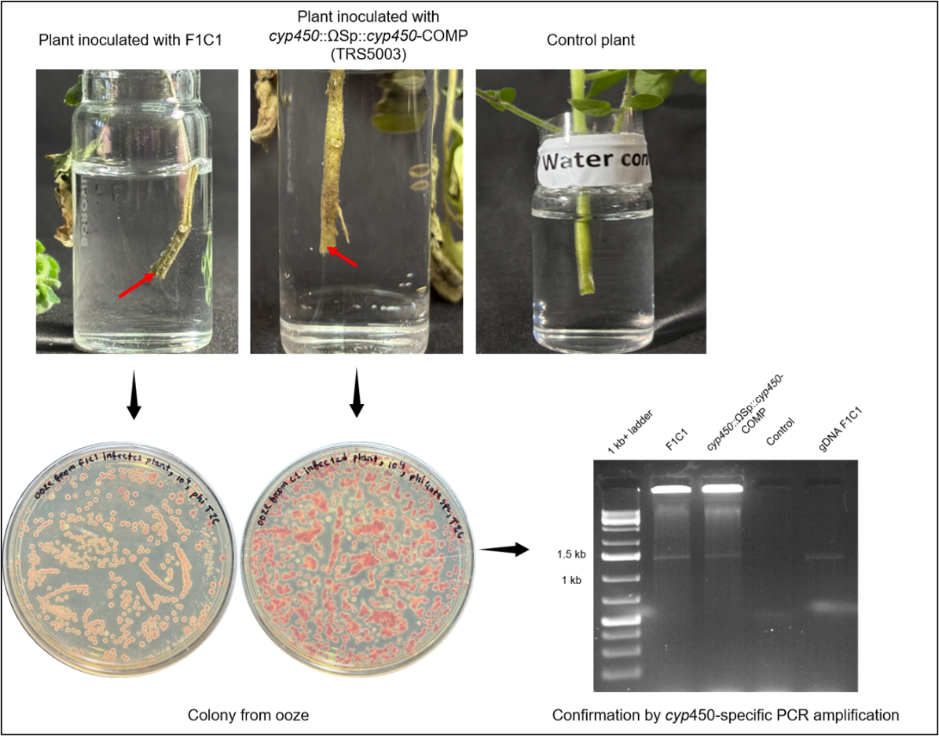

**Figure S7**

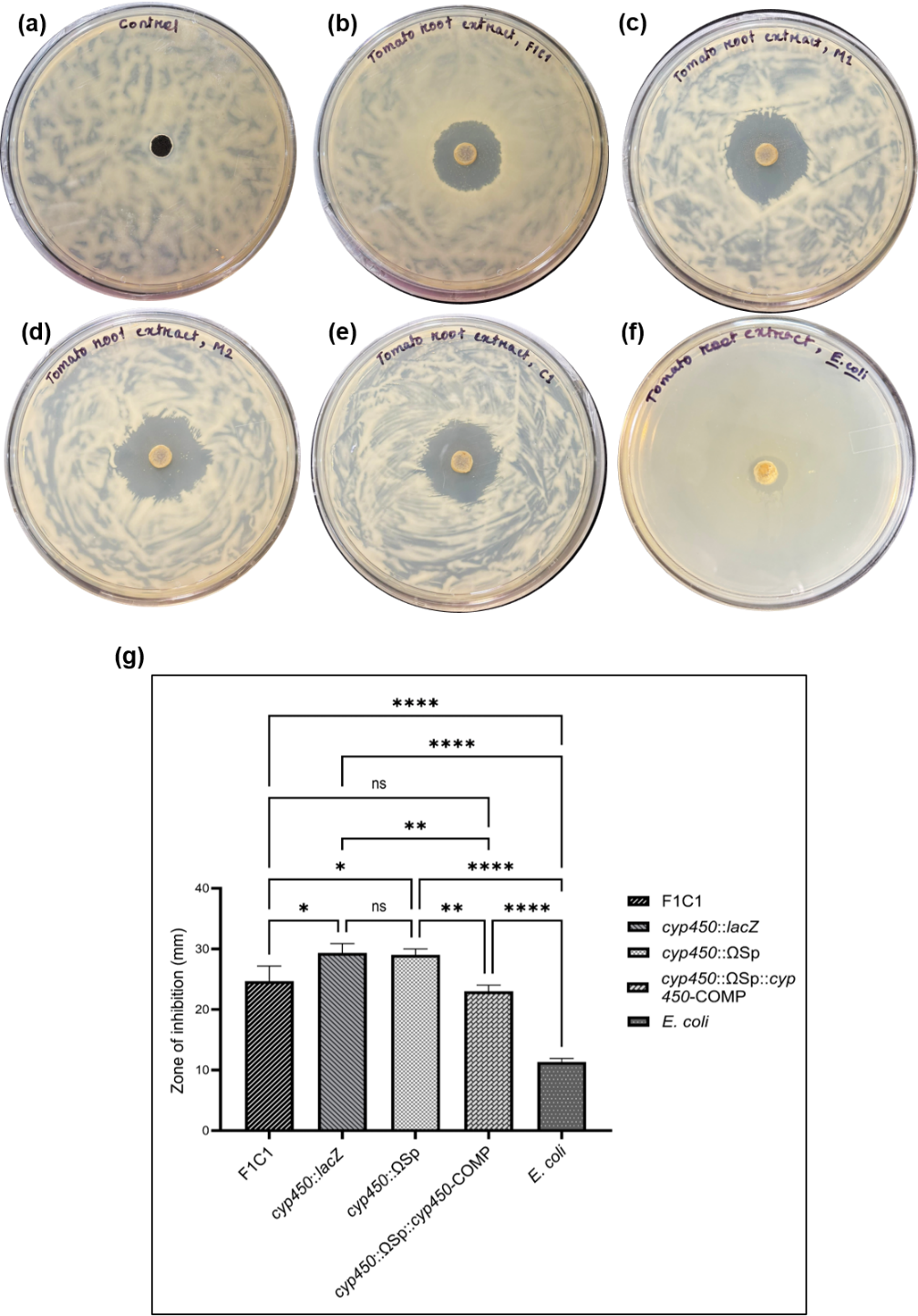

**Figure S8**

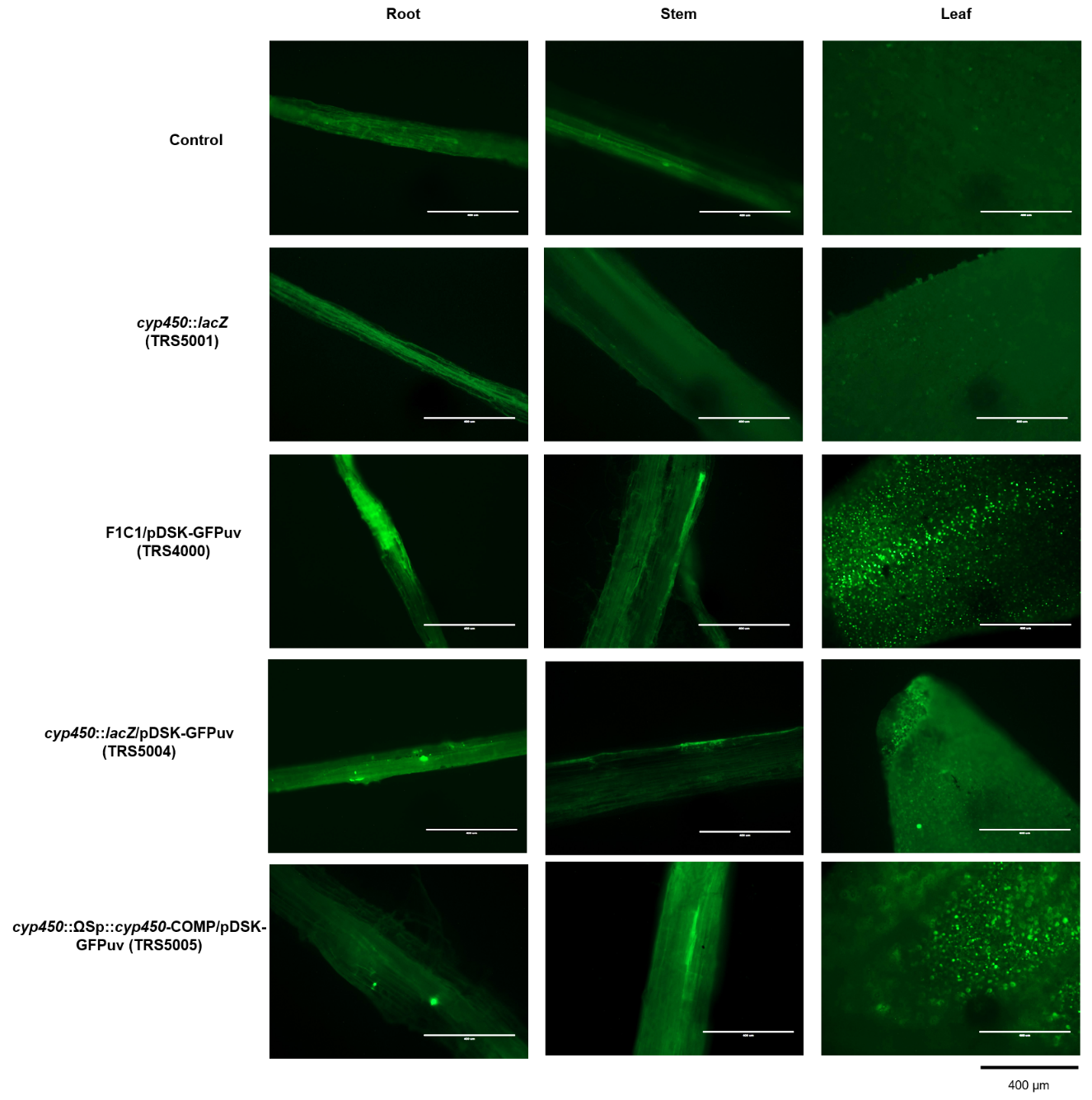

**Figure S9**

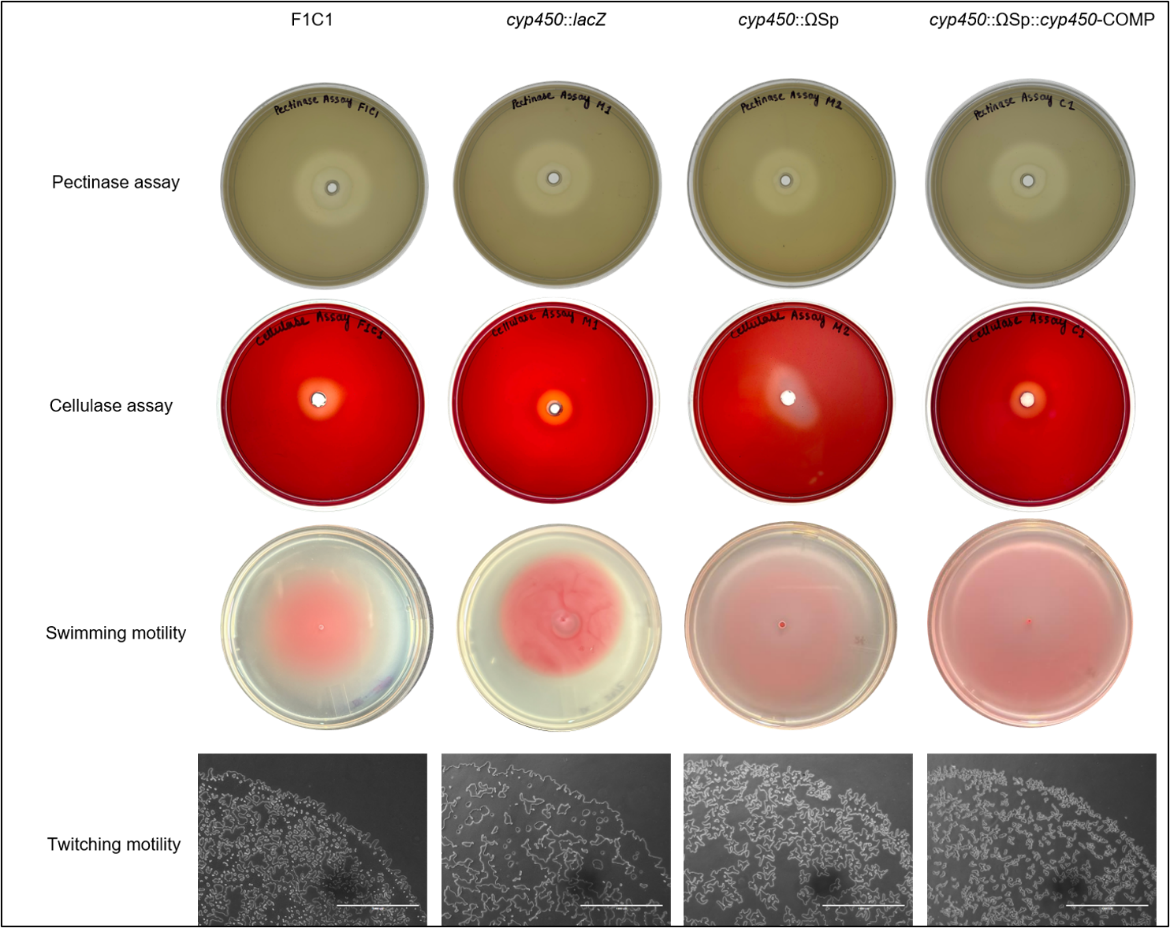

**Figure S10**
